# Sequencing and Culture-based Characterization of the Vaginal and Uterine Microbiota in Beef Cattle that Became Pregnant or Non-pregnant via Artificial Insemination

**DOI:** 10.1101/2023.07.31.551361

**Authors:** Emily M. Webb, Devin B. Holman, Kaycie N. Schmidt, Beena Pun, Kevin K. Sedivec, Jennifer L. Hurlbert, Kerri A. Bochantin, Alison K. Ward, Carl R. Dahlen, Samat Amat

**Affiliations:** Department of Microbiological Sciences, North Dakota State University, Fargo, ND, 58108, USA; Lacombe Research and Development Centre, Agriculture and Agri-Food Canada, Lacombe, AB, Canada; Central Grasslands Research Extension Center, North Dakota State University, Streeter, ND, United States; Department of Animal Sciences, and Center for Nutrition and Pregnancy, North Dakota State University, Fargo, ND, 58102, USA

**Keywords:** vaginal microbiota, uterine microbiota, bovine, 16S rRNA gene sequencing, culturing, pregnancy, artificial insemination, antimicrobial resistance

## Abstract

In this study, we evaluated the vaginal and uterine microbiota between beef cattle that became pregnant via artificial insemination (AI) and those that did not to identify microbial signature associated with pregnancy. We also characterized the culturable fraction of these microbiota using extensive culturing and screened some vaginal and uterine bacterial isolates for their antimicrobial resistance. For this, vaginal and uterine swabs from two cohorts of Angus-crossbred cattle: mature cows (vaginal and uterine; 27 open and 31 pregnant) and heifers (vaginal; 26 open and 33 pregnant) that were collected before AI were processed for microbiota assessment using 16S rRNA gene sequencing and culturing. Twenty-nine vaginal and uterine bacterial isolates were screened for resistance against 41 antibiotics. Sequencing results revealed 11 taxa that were more abundant in the vaginal samples from non-pregnant heifers compared to pregnant heifers. No differentially abundant taxa were detected in the vaginal samples from pregnant versus non-pregnant cows. Pregnant cows had a distinct uterine microbiota community structure (*P* = 0.008) and interaction network structure compared to non-pregnant cows. Twenty-eight differentially abundant uterine taxa were observed between the two groups. Community structure and diversity were different between the cow vagina and uterus. A total of 733 bacterial isolates were recovered from vaginal (512) and uterine (221) swabs under aerobic (83 different species) and anaerobic (69 species) culturing. Among these isolates were pathogenic species and those mostly susceptible to tested antibiotics. Overall, our results indicate that pregnancy-associated taxonomic signatures are present in the bovine uterine and vaginal microbiota.

**Importance:** Emerging evidence suggests that microbiome-targeted approaches may provide a novel opportunity to reduce the incidence of reproductive failures in cattle. To develop such microbiome-based strategies, one of the first logical steps is to identify reproductive microbiome features related to fertility, and isolate the pregnancy associated microbial species for developing a future bacterial consortium that could be administered before breeding to enhance pregnancy outcomes. Here, we characterized the vaginal and uterine microbiota in beef cattle that became pregnant or not via AI and identified some microbiota features associated with pregnancy. We compared similarities between vaginal and uterine microbiota, and between heifers and cows. Using extensive culturing, we provided new insights on the culturable fraction of the vaginal and uterine microbiota, and their antimicrobial resistance. Overall, our findings will serve as an important basis for future research aimed at harnessing the vaginal and uterine microbiome for improved cattle fertility.

## Introduction

Reproductive failure is a major factor in the beef and dairy cattle industry, resulting in significant and negative management and economic impacts in spite of recent advances in artificial insemination (AI), genetic selection, and improved nutrition and management of cattle over the past several decades (Ayalon, 1978; Sheldon and Dobson, 2003; Reese et al., 2020). A recent meta-analysis stated that nearly 48% of beef cow experience loss of pregnancy within the first 30 days of gestation after a single insemination, and roughly 6% of pregnancy loss occurs during the remaining gestation period (Reese et al., 2020). Therefore, improving fertility and reducing pregnancy loss in beef cattle is very important in maintaining sustainable beef production.

In recent years, numerous studies have outlined the impact that the reproductive tract microbiome may have on reproductive efficiency in humans and other vertebrate animal models, suggesting that novel, microbiome-targeted methods may reduce the incidence of reproductive failures (Koedooder et al., 2019b; Rowe et al., 2020; Tomaiuolo et al., 2020). The vaginal microbiome and its effects on fertility in women has been of particular interest in recent years (Koedooder et al., 2019b; Zhao et al., 2020). High-throughput sequencing-based studies have identified distinctive vaginal microbiota states in fertile and infertile women, and some key bacteria seem to show a species and/or strain specific effect. Zhao et al. (2020) observed that women suffering from infertility exhibited reduced vaginal microbial diversity (the number and relative abundance of microbial taxa) and richness (the number of microbial taxa present in a given environment; Hong et al., 2006) compared to fertile women. Another research group was able to predict pregnancy outcomes, with relatively high accuracy, based on the vaginal microbiota composition of 192 women, identifying a subgroup of women who were less likely to become pregnant based on their vaginal microbiota profile (Koedooder et al., 2019). These studies suggest that the vaginal microbiota is not only associated with vaginal health, but that there is a need for further research on the role that particular bacterial species play in influencing reproductive outcomes.

While direct evidence is still lacking, it has been suggested that the uterine microbiome may be involved in regulation of endometrial physiology, thereby influencing reproductive health, fertility, placentation, and pregnancy (Benner et al., 2018). Alteration of the endometrial microbiota characterized by a non-*Lactobacillus*-dominated microbial community has been associated with significant decreases in implantation rate, pregnancy establishment, and live birth weight in humans (Moreno et al., 2016). One proposed mechanism through which dysbiosis of the uterine microbiota exerts a negative effect on successful implantation is by a microbiota-triggered inflammatory response in the endometrium that ultimately interferes with the adherence of the blastocyst to the endometrial mucosa (Bardos et al., 2019). The perturbation of the uterine microbiota may negatively affect conception rate and increase pregnancy loss. Thus, maintaining homeostasis of the uterine microbiome is important in female reproductive health and fertility.

Although culture-independent techniques including 16S rRNA gene sequencing are often used to study the composition and structure of the microbial communities, they lack the ability to provide higher taxonomic resolution (species), or information on cell viability or phenotype (Koedooder et al., 2019a). Inclusion of culture-based methods along with sequencing can therefore yield a more complete understanding of microbial communities. For example, a combination of culturing and sequencing is often used in clinical settings to diagnose complex infections such as chronic endometriosis (Moreno et al., 2018). This approach is likely necessary to develop microbiome-targeted strategies to improve female fertility.

Antimicrobial resistance (AMR) is a global One Health issue that has significant implications for both human and animal health. In 2019, four million human deaths were estimated to be associated with AMR and 1.3 million of these deaths were caused by antibiotic resistant bacteria (Murray et al., 2022).The livestock industry is facing similar challenges and may suffer significant AMR-associated economic and animal welfare losses with an estimated 11% decrease in livestock production output by 2050 if current AMR trends persist (Dadgostar, 2019). The emergence and spread of AMR in zoonotic and bovine pathogens is one of the greatest challenges facing the modern beef cattle industry (Cameron and McAllister, 2016). Surveillance of AMR in microbiomes residing within the gastrointestinal (Cameron and McAllister, 2016) and respiratory (Andrés-Lasheras et al., 2021; Nobrega et al., 2021) tracts of beef cattle have been relatively well documented, and efforts to develop antimicrobial alternatives are underway (Cameron and McAllister, 2019). Antimicrobial resistance in commensal microorganisms present in the reproductive tract of female cattle has been poorly characterized despite the uterus and vagina having a very intimate relationship with the fetus and the potential role that the female tract has in the transfer of resistant bacteria from mother to offspring.

Taken together, the present study was conducted to 1) characterize the vaginal and uterine microbiota between beef cattle that became pregnant via AI and those that did not; 2) identify differentially abundant taxa between pregnant and non-pregnant cattle; 3) characterize the culturable fraction of the vaginal and uterine microbiota using extensive culturing; and 4) investigate AMR in these vaginal and uterine bacterial isolates. In addition, similarities between vaginal and uterine microbiota, and between heifers and cows (nulli-, primi-, and multi-parous) were evaluated.

## Materials and Methods

All experimental procedures involving cattle were approved by the North Dakota State University Institutional Animal Care and Use Committee (IACUC protocol# A21061 and IACUC# 21049 for cows and heifers, respectively).

### Animal husbandry and experimental design

Two cohorts of Angus based cross-bred female cattle: cows (N=100) and heifers (N = 72) were selected for collection of vaginal (cows and heifers) and uterine (cows only) swabs 2 days prior (heifers) or at the time of artificial insemination (AI) (cows). The cattle were synchronized with a 7-day Co-Synch + CIDR protocol for fixed-time AI (Lamb et al., 2010). Pregnancy diagnosis via ultrasound was performed 35 days after AI. Cows had free access to grazing pastures beginning 45 days before collection. Pasture description and species are described in the previous publication (McCarthy et al., 2023). Heifers were managed on a diet containing alfalfa/grass hay (60% of diet, dry matter basis), ground corn (31%), and corn silage (9%). These heifers were sourced from the same farm where the cows were raised, but, at the time of sample collections, they were raised in the Animal Nutrition and Physiology Center (ANPC) at North Dakota State University (Fargo, ND) and were individually fed using an electronic head-gate facility (American Calan; Northwood, NH)

### Vaginal and uterine swab collection

#### Vaginal swab collection

Vaginal swabs were collected from both cows (N = 100) at the time of AI and heifers (N = 72) 2 days before AI using the method described previously (Amat et al., 2021). For vaginal swab collection, the vulva was thoroughly cleaned with 70% ethanol and a paper towel. The labia majora was then held open allowing the passage of a swab into the vagina (15 cm, sterile cotton tipped applicators with aerated tip protector; Puritan). When the swab tip reached the midpoint of the vaginal cavity, it was swirled four times, making consistent contact with the vaginal wall, then carefully withdrawn to minimize contamination. The swabs were immediately placed into sterile Whirl-Pak bags (Uline, Pleasant Prairie, WI. USA) placed on ice and transported to the lab.

#### Uterine swab collection

Immediately after vaginal swab collection from each cow, uterine swab sampling was performed using a double guarded culture swab (71 cm in length, Reproduction Provisions L.L.C., Walworth, WI, USA). The double guarded culture swab was guided via rectal palpation through the vagina to the cervix, then the inner plastic portion was threaded through the cervix and into the uterine body. Once in the uterine body, the swab tip was extended through the inner plastic sheath. Gentle pressure was applied by pinching the uterine body to the swab via rectal palpation, then the swab was rotated 3 times. Once the sample was collected, the swab was retracted into the inner and outer plastic sheaths and removed from the cow. The tip of the swab was then cut and placed in a 2-ml tube and placed on ice and transported to the lab. Of note, both vaginal and uterine swabs from the cows were collected simultaneously prior to insemination, and these samples were collected from all 100 cows within 4 h by same personnels.

At the lab, the uterine and vaginal swabs were transferred into 1 ml brain heart infusion (BHI) broth (Hardy Diagnostics, Santa Maria, CA, USA) containing 20% glycerol (Fisher Scientific, Fair Lawn, NJ, USA) and stored at -80°C for genomic DNA extraction and culturing.

### Metagenomic DNA extraction

The DNeasy Blood and Tissue Kit (Qiagen Inc., Hilden, Germany) was used to extract genomic DNA from the vaginal and uterine swab samples. Prior to extraction, the samples were thawed, and then thoroughly vortexed before transferring 500 µl of the BHI+20% glycerol containing a swab to a new 2-ml screw-cap tube. This tube was then centrifuged at 20,000 × *g* for 10 min at 4°C, and the supernatant was carefully discarded to avoid disturbing the pellet. Using a sterile scalpel and/or forceps, the cotton tip was removed from the swab handle and added to the corresponding cell pellet. Next, 350 µl of enzymatic lysis buffer (20 mM Tris.Cl [pH 8], 2 mM sodium ethylenediaminetetraacetic acid [EDTA], 1.2% Triton X-100, lysozyme [100 mg/ml], and mutanolysin [25,000 U/ml]) was added to each tube before vortexing to dissolve the pellet and submerge the cotton. The tubes were then incubated for 1 h at 37°C with agitation at 800 rpm (Amat et al., 2021). The Dneasy Blood and Tissue Kit was then used as described in the manufacturer’s protocol with the addition of two rounds of bead beating at 6.0 m/s for 40 seconds using FastPrep-24 bead beater (MP Biomedicals, Irvine, CA, USA) after the proteinase K incubation. The DNA was eluted with 50 µl of pre-warmed elution buffer and the quantity and quality of the extracted DNA was measured using a NanoDrop ND-1000 spectrophotometer and PicoGreen assay. The DNA was stored at – 20 °C until 16S rRNA gene library preparation and sequencing.

### 16S rRNA gene sequencing and analysis

The V3-V4 hypervariable regions of the 16S rRNA gene were amplified as described previously (Amat et al., 2021; Winders et al., 2023). Briefly, Phusion High-Fidelity PCR Master Mix (New England Biolabs, Ipswich, MA, USA) was used for all PCR steps. The DNA fragment of interest was excised from a 2% agarose gel and purified with QIAquick Gel Extraction Kit (Qiagen Inc.). The NEBNext Ultra DNA Library Prep Kit (New England Biolabs) for Illumina was used for sequencing library preparation following the manufacturer recommendations. A Qubit 2.0 Fluorometer (Thermo Scientific, Waltham, MA, USA) and Agilent Bioanalyzer 2100 system were used to assess library quality. The 16S rRNA gene libraries were then sequenced on a NovaSeq 6000 instrument with a SP flow cell (2 × 250 bp) (Illumina Inc., San Diego, CA, USA).

The DADA2 v. 1.18 package (Callahan et al., 2016) in R. 4.0.3 was used to process the 16S rRNA gene sequences. The forward reads were truncated at 225 bp and the reverse reads at 220 bp. The reads were merged, chimeric sequences removed, and taxonomy assigned to each merged sequence (amplicon sequence variant [ASV]), using the naïve Bayesian RDP classifier (Wang et al., 2007) and the SILVA SSU database release 138.1 (Quast et al., 2013). The ASVs found predominantly in negative extraction control samples that were likely contaminants and ASVs classified as chloroplasts, mitochondria, or eukaryotic were removed prior to analysis. Phyloseq 1.34.0 (McMurdie and Holmes, 2013) and vegan 2.5-7 (Oksanen et al., 2013) were used to calculate the number of ASVs per sample (richness), the Shannon and inverse Simpson’ diversity indices, and Bray-Curtis dissimilarities in R. To account for uneven sequence depths, samples were randomly subsampled to 14,500 sequencing reads for the vaginal and uterine samples, prior to the calculation of Bray-Curtis dissimilarities.

### Isolation of bacteria from vaginal and uterine swab samples

#### Source of swabs used for culturing

Swabs collected from a 2021 cohort of cows (both vaginal and uterine swabs) and heifers (vaginal only) that were used for 16S rRNA gene sequencing were also processed for culturing. Bacteria from vaginal (n = 12) and uterine (n = 30) swabs collected from a similar cohort of cows and heifers in the following year (2022) were also cultured to increase the isolation of more diverse bacterial species from the female reproductive tract. Cryopreserved vaginal and uterine swabs were subjected to aerobic and anaerobic culturing using different growth media as described below.

#### Aerobic culturing

A 50-µl aliquot of undiluted BHI + 20% glycerol stock from uterine and vaginal swab samples was spread onto De Man, Rogosa, and Sharpe (MRS) agar (Hardy Diagnostics, Santa Maria, CA) and Columbia blood agar plates supplemented with 5% sheep’s blood (CB) (Becton, Dickinson and Company, Sparks, MD, USA). Similarly, 50 µl of undiluted vaginal sample was spread on MRS agar, while 50 µl of vaginal sample diluted 1:10 in phosphate buffered saline (PBS) (Mediatech Inc., Manasas, VA, USA) was spread on CB agar. The CB agar plates were incubated at 37°C in 5% CO_2_ and the MRS agar plates at 37°C in 10% CO_2_ for up to 48 h. Up to eight colonies with distinctive morphologies on each plate were then subcultured onto its respective agar and incubated under the same conditions as above. After visually assessing the purity of each isolate, a disposable 100 µl loop was used to transfer each isolate into 100 µl Tris-EDTA (TE) (Quality Biological Inc., Gaithersburg, MD, USA) stock and 1 ml of 20% (v/v) glycerol containing MRS or BHI broth depending on what agar the isolate was grown on. The TE stocks were stored at -20°C for genomic DNA extraction, while the MRS (MRSg) and BHI glycerol (BHIg) stocks were stored at -80°C.

#### Anaerobic culturing

Anaerobic culturing was performed in an anaerobic chamber (Type B, Vinyl, Coy Laboratory Products Inc., Grass Lake, MI, USA) supplied with a gas mixture containing 90% N_2_, 5% CO_2_, and 5% H_2_. A subset of vaginal and uterine samples from cows were diluted 1:2 with PBS, and 100 µl was plated onto MRS agar while 50 µl was plated on blood and Wilkins Chalgren (WC) (HiMedia Laboratories, Mumbai, India) agars. These agar plates were incubated at 37°C for up to 72 h. Colonies with a unique morphology were selected and subcultured onto a CB agar plate and incubated at 37°C for up to 72 h. A loopful of each isolate was transferred to 100-µl TE and 1-ml BHI glycerol stock and then stored at -20°C or -80°C, respectively, until used for DNA extraction or culturing.

### Identification of uterine and vaginal isolates

Bacterial strains were identified using amplification and sequencing of near full length 16S rRNA gene. For this, genomic DNA was extracted from all MRS (aerobic: n = 122; anaerobic n = 104), CB (n = 241), Blood (n = 135), and WC (anaerobic: n = 131) isolates using the Quick-DNA Fungal/Bacterial Miniprep Kit (Zymo Research, Irvine, CA, USA) according to manufacturer’s instructions with the modifications outlined in our previous publication (Webb et al., 2023).

The near-full length 16S rRNA gene was amplified via PCR using the universal primers 27F (5’-AGAGTTTGATCMTGGCTCAG -3’) and 1492R (5’-TACGGYTACCTTGTTACGACTT -3’) as described previously (Amat et al., 2019; Webb et al., 2023). Each PCR consisted of 20-µl iQ Supermix (Bio-Rad Laboratories, Inc., Hercules, CA, USA), 1 µl of each primer (10 µM), and 2 µl of isolate DNA for a total volume of 40 µl per reaction. The PCR conditions were as follows: an initial denaturation of 95°C for 5 min; 35 cycles of 95°C for 45 s, 50°C for 30 s, 72°C for 2 min; and a final extension at 72°C for 5 min. The PCR reactions were performed using an Eppendorf Mastercycler (Eppendorf, Hamburg, Germany). A 1% (w/v) agarose gel was used to visualize the PCR products and amplicons were sent to MCLAB (San Francisco, CA, USA) for Sanger sequencing. The 16S rRNA gene sequences were identified using the Basic Local Alignment Search Tool (BLAST) and the non-redundant NCBI nucleotide database.

### Evaluation of antimicrobial susceptibilities of selected vaginal and uterine isolates

To gain insight into the AMR in commensal bacteria residing within the female bovine reproductive tract, a total of 29 bacterial isolates (10 Gram-positive and 19 Gram-negative) were selected and antimicrobial susceptibility testing (AST) was performed. The isolate selection criteria for inclusion were based on isolation frequency from the vaginal and uterine swabs, the ability to culture the isolate aerobically as required by the AST testing method, and most importantly, the availability of the antibiotic breakpoints. Antimicrobial susceptibility testing was performed on fresh colonies at the NDSU Veterinary Diagnostic Laboratory (Fargo, ND) as described previously (Webb et al., 2023). The minimum inhibitory concentrations (MICs) of 41 antibiotics were determined by microdilution (Sensititre; Thermo Fisher Scientific, Nepean, ON, Canada) using commercially available panels (companion animal Gram-positive [COMPGP1F], equine [EQUIN1F], bovine [BOPO7F] Trek Diagnostic Systems, Cleveland, OH, USA) (Supplementary Table S2, Table S3). The AST set-up and procedure were performed as suggested for these panels. The Sensititre AIM delivery system was used to inoculate to the 96-well AST plates and the plates were incubated aerobically at 37 °C for 24 h. After incubation, the plates were evaluated with a BIOMIC V3 (Giles Scientific USA; Santa Barbara, CA, USA). Quality control was performed on the microdilution plates as recommended by the Clinical and Laboratory Standards Institute (CLSI) VET01S standards (CLSI, 2021). Breakpoints for *Corynebacterium* isolates were determined using the criteria provided by CLSI document M45 (CLSI, 2016). Breakpoints for all other isolates were determined using CLSI M100 (Supplementary Table 2 and 3) (CLSI, 2022).

### Statistical analysis

Bray-Curtis dissimilarities and PERMANOVA using the adonis2 function in vegan in R were used to assess the vaginal and uterine microbial community structure in pregnant vs. open (non-pregnant) cattle. To identify pregnancy-associated taxonomic signatures, differentially abundant taxa in the vaginal and uterine microbiota between pregnant and non-pregnant cows and heifers were identified using the DESeq2 package. Only those ASVs found in at least 25% of all samples analyzed were included in the DESeq2 analyses.

The number of ASVs (microbial richness), diversity indices, relative abundance of the most relatively abundant phyla and genera between pregnant and open groups were compared using the t-test or the Mann-Whitney U test depending on the normality the data using SAS (ver. 9.4, SAS Institute Inc., Cary, NC, USA). The Shapiro-Wilk test was used to determine whether a dataset follows a normal distribution. Significance was considered at *P* < 0.05.

Ecological network modeling was performed to compare the interaction network structure among all observed genera in uterine microbiota between non-pregnant and pregnant cows using the generalized Lotka-Volterra models (gLVMs) by coupling biomass estimation and model inference with an expectation-maximization algorithm (BEEM) as described by Li et al.(Li et al., 2019). The interaction network models (open and pregnant groups) created by BEEM-statistic was visualized by using the graph package of R (Amat et al., 2023).

## Results

### 16S rRNA gene sequencing overview

After processing and quality filtering, the average number of sequences per sample from the vaginal and uterine samples (n = 119) obtained from cows, and heifers (vaginal swabs only, n = 60) were 58, 928 ± 23,109 (SD), and 85,343 ± 23,422, respectively. From these sequences, a total of 1,114 unique genera were identified and classified into 41 phyla: 34 bacterial (accounting for 99.7% of total sequences) and 7 archaeal (0.23%) phyla.

### The vaginal microbiota in heifers that became pregnant or not via AI

Community structure of the vaginal microbiota did not differ between the pregnant and non-pregnant heifers at the time of AI (R^2^ = 0.017; *P* = 0.04) (Fig.1A). Pregnant heifers had similar microbial richness (observed ASVs) (Fig. 1B), and diversity (Shannon and inverse Simpson diversity indices; Fig. 1C, D) compared to non-pregnant heifers (*P* > 0.05).

**Figure 1.**
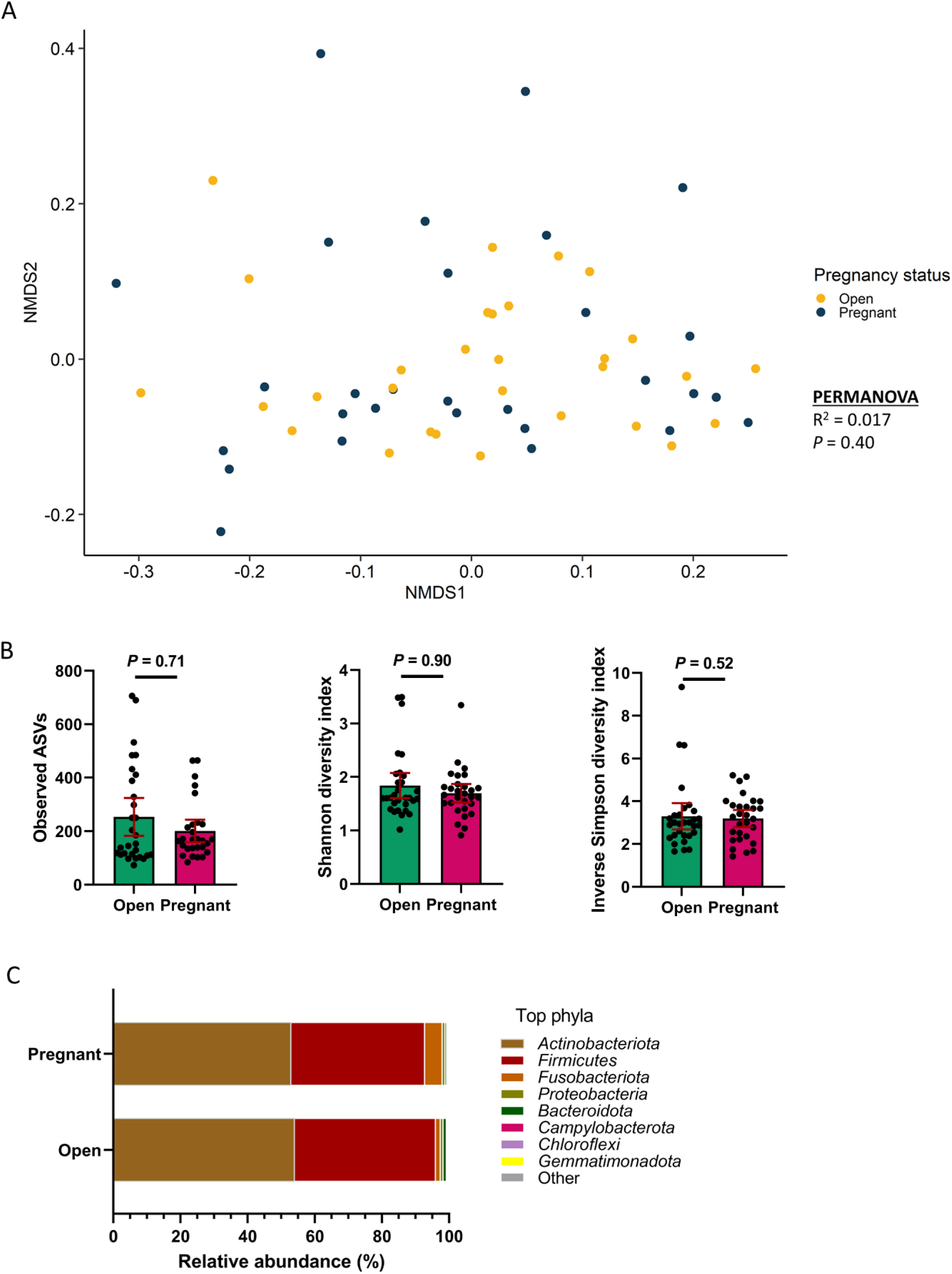
Beta and alpha diversity, and composition (phylum level) of the vaginal microbiota of non-pregnant (open, n = 33) and pregnant heifers (n = 26). (**A**) Nonmetric multidimensional scaling (NMDS) plots of the Bray–Curtis dissimilarities, (**B**) number of observed amplicon sequence variants (ASVs), and Shannon and inverse Simpson’ diversity index, (**C**) percent relative abundance of the eight most relatively abundant phyla.

No differences were observed in relative abundance of top 7 relatively most abundant bacterial phyla (Fig. 1D) and top 20 genera (Table 1) in the vaginal microbiota between the two group heifers (*P* > 0.05). However, the DESeq2 identified 11 ASVs that were significantly enriched in the vaginas of heifers that failed to become pregnant via AI compared to heifers that were successfully impregnated (*P* < 0.10) (Table 2). These ASVs were all within the phylum *Firmicutes*, although only four were classified at genus level. The ASVs classified at genus level were within the family *Lachnospiraceae* (*dorea*, [ASV19], *Coprococcus* [ASV59] and UCG-010 [ASV30]), *Oscillospiraceae* (UCG-005 [ASV27 and 112]) *Oscillibacter* [ASV132]) and *Butyricicoccaceae* (UCG-009 [ASV86]).

**Table 1.**
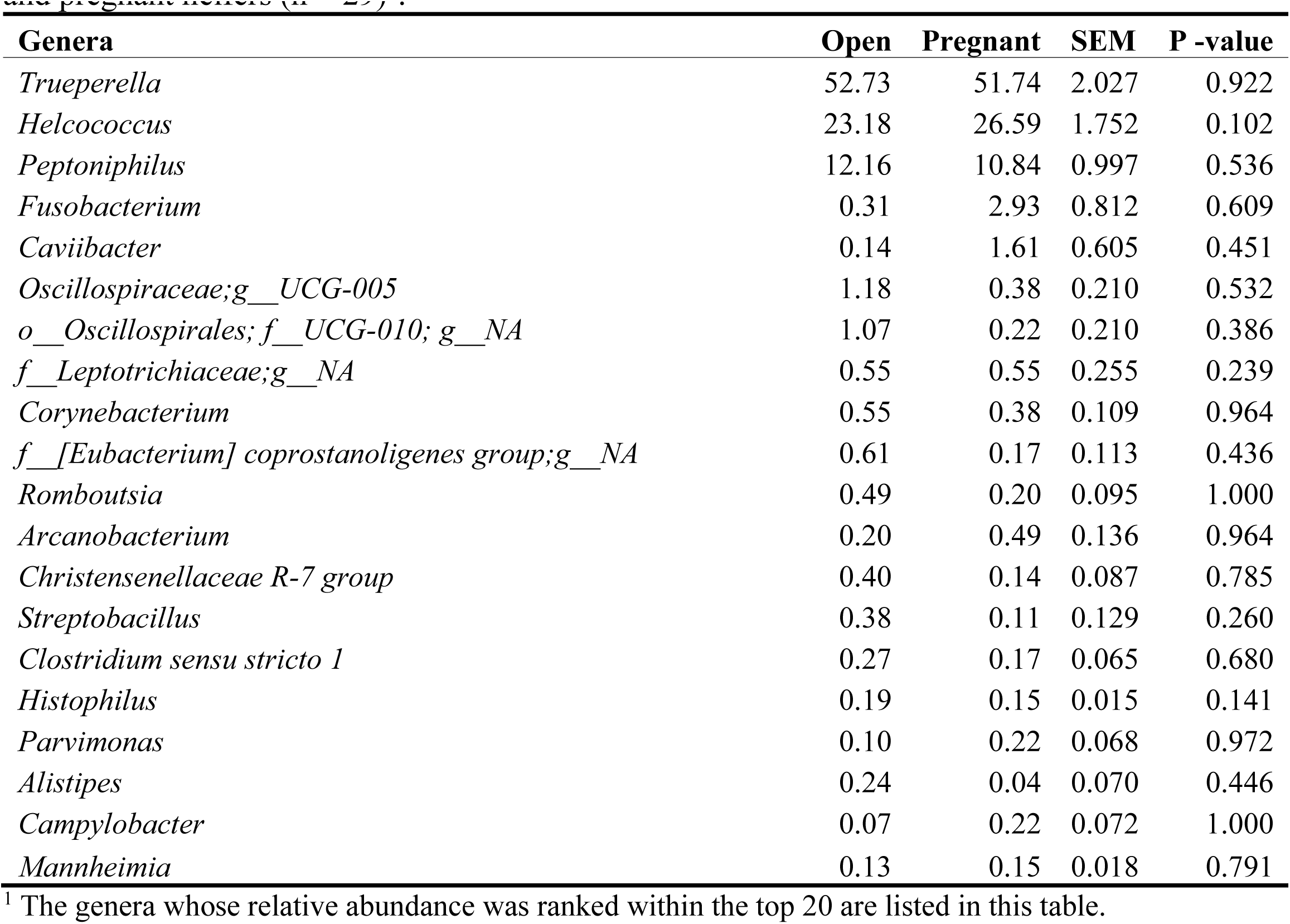
Top 20 most relatively abundant bacterial genera in vaginal microbiota of open (n = 31) and pregnant heifers (n = 29)^1^.

| Genera | Open | Pregnant | SEM | P -value |
| --- | --- | --- | --- | --- |
| *Trueperella* | 52.73 | 51.74 | 2.027 | 0.922 |
| *Helcococcus* | 23.18 | 26.59 | 1.752 | 0.102 |
| *Peptoniphilus* | 12.16 | 10.84 | 0.997 | 0.536 |
| *Fusobacterium* | 0.31 | 2.93 | 0.812 | 0.609 |
| *Caviibacter* | 0.14 | 1.61 | 0.605 | 0.451 |
| *Oscillospiraceae*;g__UCG-005 | 1.18 | 0.38 | 0.210 | 0.532 |
| *o__Oscillospirales*; *f__UCG-010*; *g__NA* | 1.07 | 0.22 | 0.210 | 0.386 |
| *f__Leptotrichiaceae*;g__NA | 0.55 | 0.55 | 0.255 | 0.239 |
| *Corynebacterium* | 0.55 | 0.38 | 0.109 | 0.964 |
| *f__[Eubacterium] coprostanoligenes group*;g__NA | 0.61 | 0.17 | 0.113 | 0.436 |
| *Romboutsia* | 0.49 | 0.20 | 0.095 | 1.000 |
| *Arcanobacterium* | 0.20 | 0.49 | 0.136 | 0.964 |
| *Christensenellaceae R-7 group* | 0.40 | 0.14 | 0.087 | 0.785 |
| *Streptobacillus* | 0.38 | 0.11 | 0.129 | 0.260 |
| *Clostridium sensu stricto 1* | 0.27 | 0.17 | 0.065 | 0.680 |
| *Histophilus* | 0.19 | 0.15 | 0.015 | 0.141 |
| *Parvimonas* | 0.10 | 0.22 | 0.068 | 0.972 |
| *Alistipes* | 0.24 | 0.04 | 0.070 | 0.446 |
| *Campylobacter* | 0.07 | 0.22 | 0.072 | 1.000 |
| *Mannheimia* | 0.13 | 0.15 | 0.018 | 0.791 |

<sup>1</sup> The genera whose relative abundance was ranked within the top 20 are listed in this table.

**Table 2.**
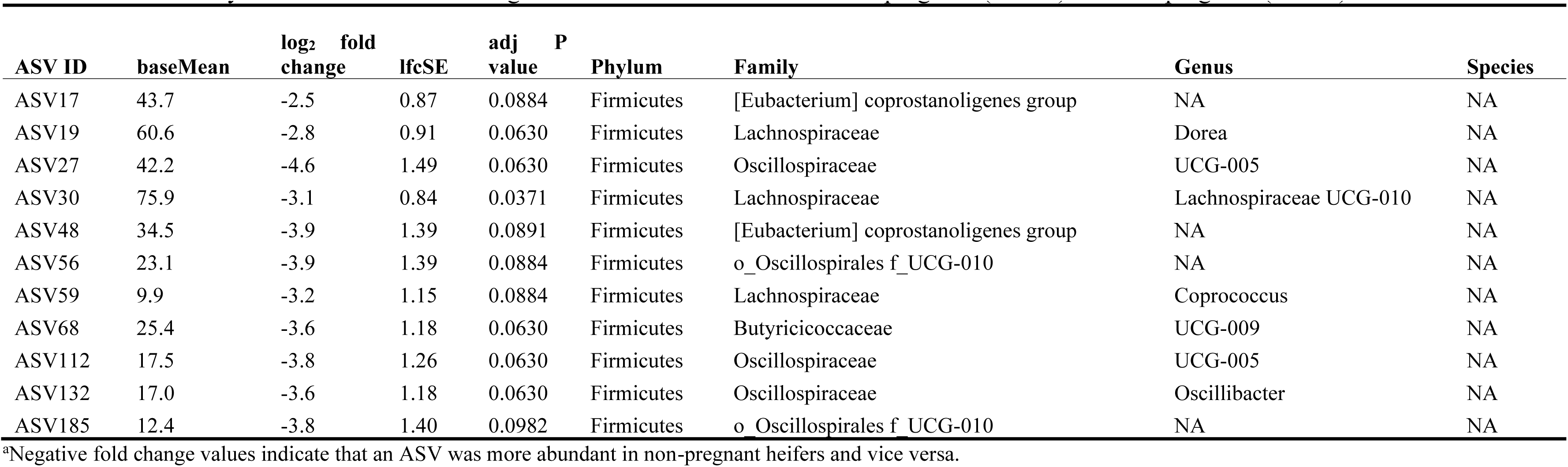
Differentially abundant ASVs in the vaginal microbiota of heifers between pregnant (n = 33) and non-pregnant (n = 26)^a^.

| ASV ID | baseMean | log<sub>2</sub> fold change | lfcSE | adj P value | Phylum | Family | Genus | Species |
| --- | --- | --- | --- | --- | --- | --- | --- | --- |
| ASV17 | 43.7 | -2.5 | 0.87 | 0.0884 | Firmicutes | [Eubacterium] coprostanoligenes group | NA | NA |
| ASV19 | 60.6 | -2.8 | 0.91 | 0.0630 | Firmicutes | Lachnospiraceae | Dorea | NA |
| ASV27 | 42.2 | -4.6 | 1.49 | 0.0630 | Firmicutes | Oscillospiraceae | UCG-005 | NA |
| ASV30 | 75.9 | -3.1 | 0.84 | 0.0371 | Firmicutes | Lachnospiraceae | Lachnospiraceae UCG-010 | NA |
| ASV48 | 34.5 | -3.9 | 1.39 | 0.0891 | Firmicutes | [Eubacterium] coprostanoligenes group | NA | NA |
| ASV56 | 23.1 | -3.9 | 1.39 | 0.0884 | Firmicutes | o_Oscillospirales f_UCG-010 | NA | NA |
| ASV59 | 9.9 | -3.2 | 1.15 | 0.0884 | Firmicutes | Lachnospiraceae | Coprococcus | NA |
| ASV68 | 25.4 | -3.6 | 1.18 | 0.0630 | Firmicutes | Butyricicoccaceae | UCG-009 | NA |
| ASV112 | 17.5 | -3.8 | 1.26 | 0.0630 | Firmicutes | Oscillospiraceae | UCG-005 | NA |
| ASV132 | 17.0 | -3.6 | 1.18 | 0.0630 | Firmicutes | Oscillospiraceae | Oscillibacter | NA |
| ASV185 | 12.4 | -3.8 | 1.40 | 0.0982 | Firmicutes | o_Oscillospirales f_UCG-010 | NA | NA |

<sup>a</sup>Negative fold change values indicate that an ASV was more abundant in non-pregnant heifers and vice versa.

### The vaginal microbiota in cows that became pregnant or not via AI

As with the heifers, the vaginal microbiota community structure was similar for pregnant and non-pregnant mature cows at the time of AI (*R^2^*= 0.020, *P* = 0.21) (Fig. 2A). Non-pregnant cows tended to have greater microbial richness (*P* = 0.054) and Shannon diversity (*P* = 0.051) compared to pregnant cows (Fig. 2B). Overall, the vaginal bacterial community in cows were predominantly colonized by the phyla Actinobacteriota and Firmicutes which accounted for almost 90% of the total community composition (Fig. 2C). Their relative abundance did not differ between pregnant and non-pregnant cows (*P* > 0.05). The third most relatively abundant phylum, Fusobacteriota, was more relatively abundant in cows (*P* < 0.05). The relative abundance of five genera differed (*P*< 0.05) between the two group cows, with *Monoglobus*, *Prevetellaceae* UCG-003, and *Anaerovoracaceae* Family XIII AD3011 being more abundant in non-pregnant cows (Table 3). However, no ASVs were differentially abundant between the two group cows (*P* > 0.1).

**Figure 2.**
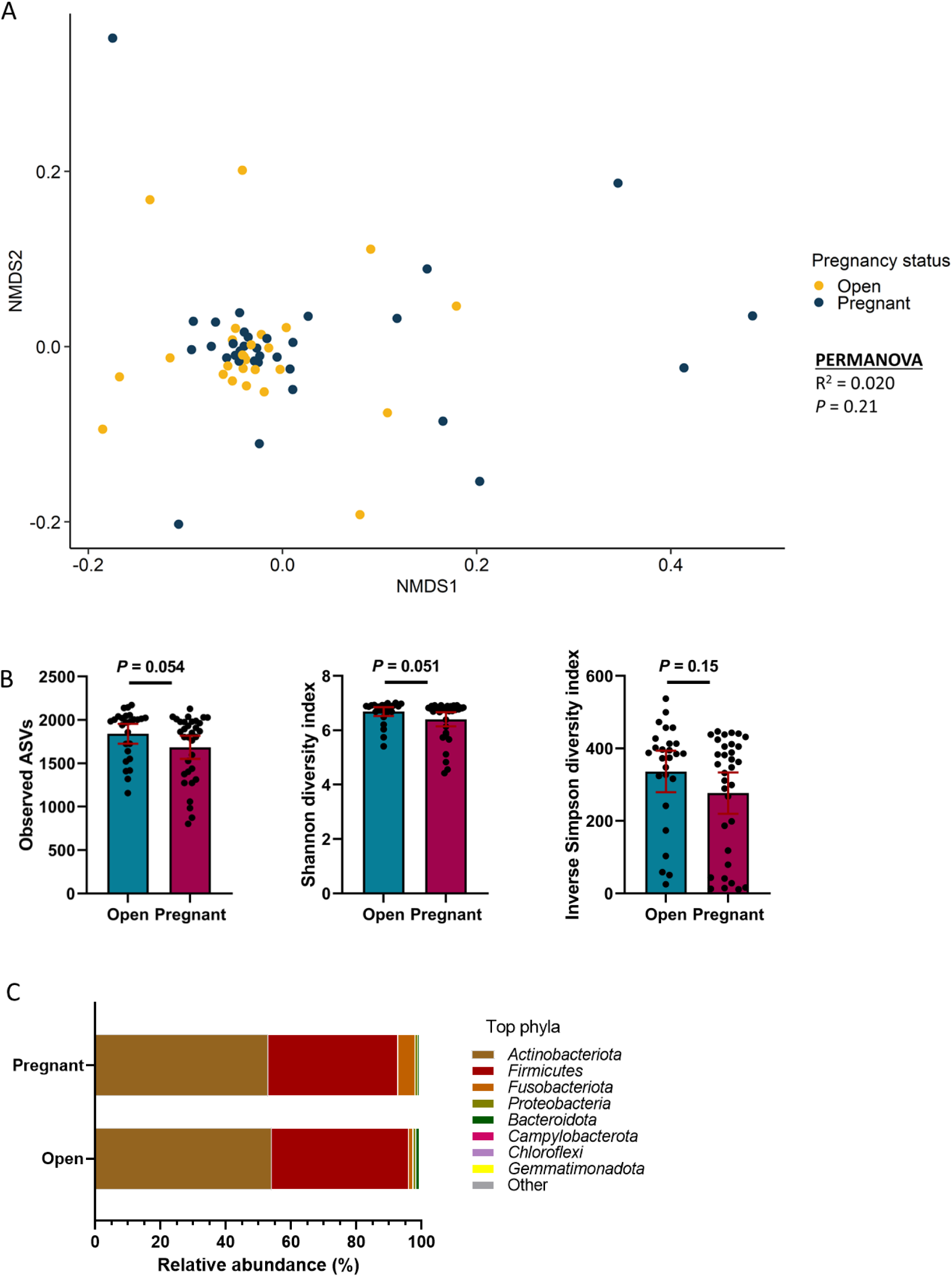
Beta and alpha diversity, and composition (phylum level) of the vaginal microbiota of non-pregnant (open, n = 31) and pregnant cows (n = 29). (**A**) Nonmetric multidimensional scaling (NMDS) plots of the Bray–Curtis dissimilarities, (**B**) number of observed amplicon sequence variants (ASVs), and Shannon and inverse Simpson’ diversity index, (**C**) percent relative abundance of the eight most relatively abundant phyla.

**Table 3.** Top 25 most relatively abundant bacterial genera in vaginal microbiota of open (n = 26) and pregnant cows (n = 33)^1^.

| Genera | Open | Pregnant | SEM | P-value |
| --- | --- | --- | --- | --- |
| *o__Oscillospirales;f__UCG-010;g__NA* | 14.52 | 13.52 | 0.423 | 0.943 |
| *f__Oscillospiraceae;g__UCG-005* | 13.95 | 13.67 | 0.313 | 0.054 |
| *f__[Eubacterium] coprostanoligenes group;g__NA* | 6.83 | 6.33 | 0.126 | 0.113 |
| *Rikenellaceae RC9 gut group* | 4.95 | 4.44 | 0.149 | 0.641 |
| *Christensenellaceae R-7 group* | 3.94 | 3.57 | 0.105 | 0.260 |
| *Alistipes* | 3.33 | 3.10 | 0.091 | 0.777 |
| *Bacteroides* | 3.25 | 3.12 | 0.095 | 0.590 |
| *Streptobacillus* | 1.70 | 4.34 | 0.876 | 0.329 |
| *Corynebacterium* | 2.85 | 1.95 | 0.472 | 0.444 |
| *Romboutsia* | 2.27 | 2.30 | 0.087 | 0.687 |
| *Monoglobus* | 2.07 | 1.83 | 0.056 | 0.033 |
| *f__Lachnospiraceae;g__NA* | 1.95 | 1.89 | 0.047 | 0.563 |
| *f__Ruminococcaceae;g__NA* | 1.92 | 1.80 | 0.049 | 0.432 |
| *f__Oscillospiraceae;g__NK4A214 group* | 1.65 | 1.56 | 0.047 | 0.755 |
| *f__Bacteroidales RF16 group;g__NA* | 1.64 | 1.54 | 0.047 | 0.959 |
| *f__Leptotrichiaceae;g__NA* | 0.47 | 2.39 | 0.584 | 0.041 |
| *o__Clostridia vadinBB60 group;f__NA;g__NA* | 1.45 | 1.37 | 0.044 | 0.799 |
| *Prevotellaceae UCG-004* | 1.35 | 1.25 | 0.043 | 0.885 |
| *f__Oscillospiraceae;g__NA* | 1.27 | 1.26 | 0.040 | 0.002 |
| *Prevotellaceae UCG-003* | 1.29 | 1.20 | 0.039 | 0.048 |
| *g__Leptotrichia* | 0.98 | 1.32 | 0.392 | 0.155 |
| *f__Butyricicoccaceae;g__UCG-009* | 1.19 | 1.15 | 0.026 | 0.873 |
| *f__Anaerovoracaceae;g__Family XIII AD3011 group* | 1.18 | 1.15 | 0.030 | 0.032 |
| *Akkermansia* | 1.13 | 1.13 | 0.108 | 0.710 |
| *o__Bacteroidales;f__NA;g__NA* | 1.06 | 0.96 | 0.036 | 0.392 |

<sup>1</sup> The genera whose relative abundance was ranked within the top 25 are listed in this table.

### The uterine microbiota in cows that became pregnant or not via AI

Pregnant cows had a distinct uterine microbiota community structure (*R^2^* = 0.032, *P* = 0.008) (Fig.3A) compared to that of non-pregnant cows. The microbial richness (observed ASVs), Shannon diversity (*P* ≥ 0.40; Fig. 3B), and inverse Simpson’s diversity index (*P* = 0.064) did not differ in the pregnant group compared to the non-pregnant group (Fig. 3B). Overall, the uterine bacterial community in cows was dominated by the phylum Firmicutes whose relative abundance accounted for almost 70% of the total 16S rRNA gene sequences (Fig. 3C). The second and third most relatively abundant phyla were Bacteroidota and Actinobacteriota. Bacteroidota abundance was elevated in non-pregnant cows (*P* < 0.05). At the genus level, among the top 25 genera listed in Table 4, 6 of them were differentially abundant (*P* < 0.05) between the two groups of cows. Four genera including *Clostridium sensu stricto 7, Fusobacterium* and the 2 genera which was not classified at genus level were enriched (*P* ≤ 0.042) in the uterus of pregnant cows. *Prevotellaceae* UCG-004 and an unclassified genus within the family *Lachnospiraceae*, however, become more abundant (*P* ≤ 0.004) in non-pregnant cows. Twenty-eight differentially abundant taxa were observed between pregnant and non-pregnant groups, with 11 of them more abundant in pregnant cows (Table 5). *Methanobrevibacter ruminantium* (archaea) and *Fusobacterium necrophorum* were among these 11 pregnancy-associated taxa.

**Figure 3.**
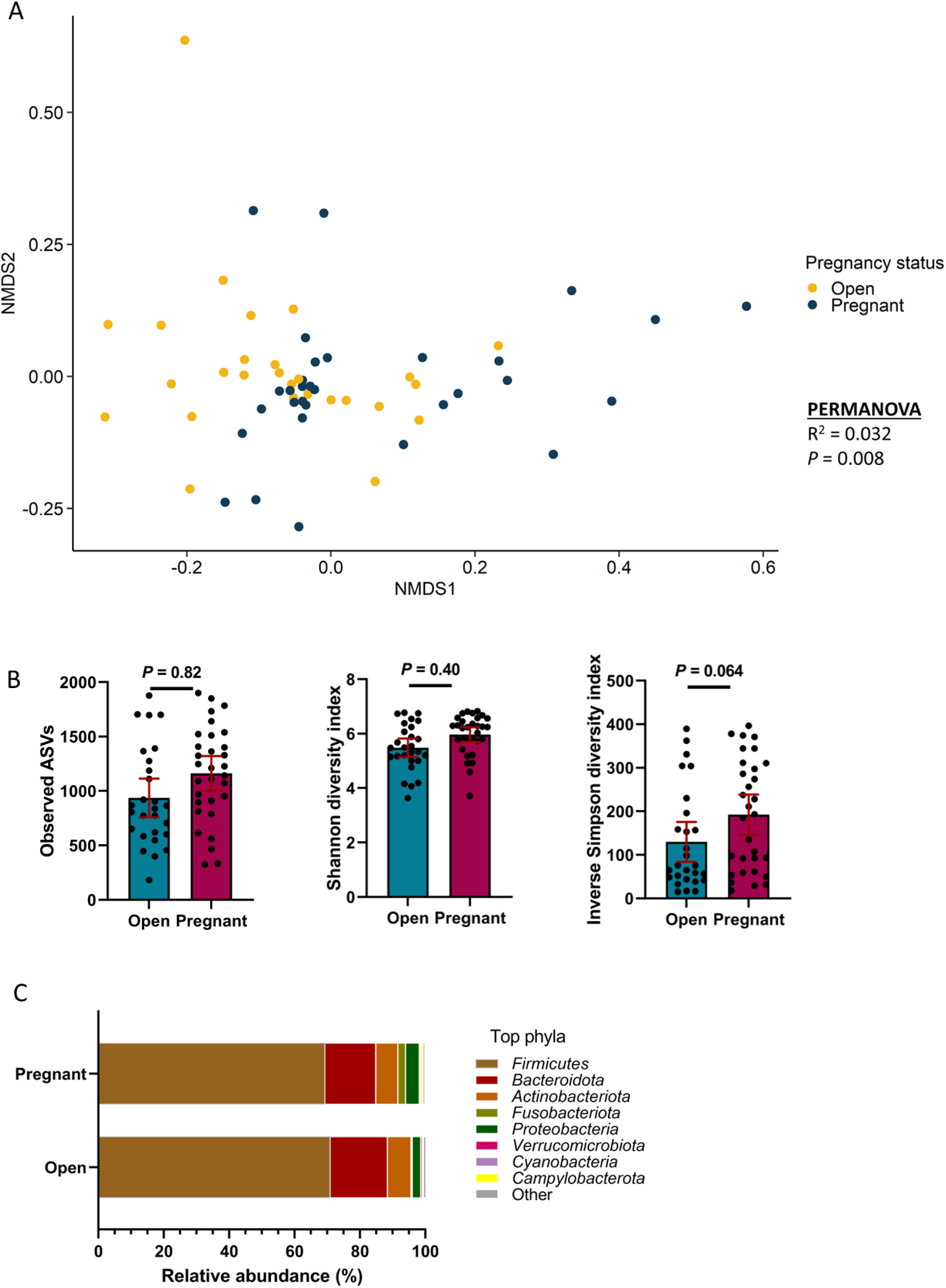
Beta and alpha diversity, and composition (phylum level) of the uterine microbiota of non-pregnant (open, n = 31) and pregnant cows (n = 29). (**A**) Nonmetric multidimensional scaling (NMDS) plots of the Bray–Curtis dissimilarities, (**B**) number of observed amplicon sequence variants (ASVs), and Shannon and inverse Simpson’ diversity index, (**C**) percent relative abundance of the eight most relatively abundant phyla.

**Table 4.** Top 25 most relatively abundant bacterial genera in the uterine microbiota of open (n = 26) and pregnant cows (n = 33)^1^.

| Genera | Open | Pregnant | SEM | P-value |
| --- | --- | --- | --- | --- |
| *o__Oscillospirales;f__UCG-010;g__NA* | 14.37 | 14.13 | 0.644 | 0.906 |
| *f__Oscillospiraceae;g__UCG-005* | 13.69 | 12.91 | 0.473 | 0.847 |
| *f__[Eubacterium] coprostanoligenes group;g__NA* | 6.86 | 6.08 | 0.275 | 0.065 |
| *Christensenellaceae R-7 group* | 3.23 | 3.40 | 0.193 | 0.443 |
| *Helcococcus* | 1.52 | 4.40 | 0.776 | 0.275 |
| *Romboutsia* | 2.99 | 2.70 | 0.222 | 0.859 |
| *Rikenellaceae RC9 gut group* | 2.74 | 2.89 | 0.213 | 0.985 |
| *Trueperella* | 1.96 | 3.27 | 0.596 | 0.321 |
| *Alistipes* | 2.29 | 2.08 | 0.131 | 0.524 |
| *Bacteroides* | 2.34 | 1.91 | 0.181 | 0.409 |
| *Clostridium sensu stricto 7* | 1.68 | 2.45 | 0.289 | 0.026 |
| *p__Proteobacteria;c__NA;o__NA;f__NA;g__NA* | 0.96 | 2.47 | 0.272 | 0.002 |
| *f__Oscillospiraceae;g__NK4A214 group* | 1.65 | 1.42 | 0.110 | 0.813 |
| *f__Lachnospiraceae;g__NA* | 1.65 | 1.42 | 0.101 | 0.004 |
| *Corynebacterium* | 1.86 | 1.21 | 0.305 | 0.353 |
| *f__Ruminococcaceae;g__NA* | 1.35 | 1.53 | 0.080 | 0.041 |
| *Prevotellaceae UCG-003* | 1.09 | 1.68 | 0.260 | 0.376 |
| *o__Clostridia vadinBB60 group;f__NA;g__NA* | 1.19 | 1.58 | 0.091 | 0.042 |
| *Monoglobus* | 1.52 | 1.18 | 0.086 | 0.557 |
| *f__Oscillospiraceae;g__NA* | 1.24 | 1.23 | 0.079 | 0.147 |
| *f__Bacteroidales RF16 group;g__NA* | 1.08 | 1.23 | 0.068 | 0.286 |
| *f__Anaerovoracaceae;g__Family XIII AD3011 group* | 1.12 | 1.08 | 0.071 | 0.792 |
| *Prevotella* | 1.33 | 0.80 | 0.307 | 0.206 |
| *Peptoniphilus* | 0.76 | 1.18 | 0.266 | 0.566 |
| *Prevotellaceae UCG-004* | 1.22 | 0.71 | 0.104 | <.0001 |
| *Fusobacterium\** | 0.005 | 0.627 | 0.1953 | 0.030 |

<sup>1</sup> The genera whose relative abundance was ranked within the top 25 are listed in this table. \* This genus whose relative abundance was ranked as 47<sup>th</sup> top genus included in this table because of its relevance in pregnancy determined by the species encompassed by this genus.

**Table 5.** Differentially abundant ASVs in the uterine microbiota of cows between pregnant (n = 31) and non-pregnant (n = 27)^a^.

| ASV ID | base Mean | log<sub>2</sub> fold change | lfcSE | adj P value | Phylum | Family | Genus | Species |
| --- | --- | --- | --- | --- | --- | --- | --- | --- |
| ASV644 | 16.7 | -5.64 | 1.33 | 0.0053 | Firmicutes | Monoglobaceae | Monoglobus | NA |
| ASV1493 | 11.2 | -4.41 | 1.34 | 0.0456 | Firmicutes | [Eubacterium] coprostanoligenes group | NA | NA |
| ASV592 | 27.9 | -4.24 | 1.36 | 0.0747 | Firmicutes | Oscillospiraceae | UCG-005 | NA |
| ASV957 | 30.8 | -4.11 | 1.15 | 0.0194 | Firmicutes | Ruminococcaceae | Faecalibacterium | prausnitzii |
| ASV612 | 10.0 | -3.96 | 0.90 | 0.0036 | Firmicutes | Christensenellaceae | Christensenellaceae R-7 group | NA |
| ASV866 | 12.1 | -3.55 | 1.20 | 0.0930 | Actinobacteriota | NA | NA | NA |
| ASV385 | 21.2 | -3.47 | 1.12 | 0.0747 | Bacteroidota | Prevotellaceae | Prevotella_7 | NA |
| ASV682 | 19.7 | -3.42 | 1.19 | 0.0970 | Firmicutes | Lachnospiraceae | [Ruminococcus] gauvreauii group | NA |
| ASV506 | 13.9 | -2.74 | 0.95 | 0.0970 | Bacteroidota | Rikenellaceae | Rikenellaceae RC9 gut group | NA |
| ASV401 | 12.9 | -2.49 | 0.81 | 0.0753 | Firmicutes | Oscillospiraceae | UCG-002 | NA |
| ASV103 | 63.7 | -2.04 | 0.52 | 0.0083 | Bacteroidota | Prevotellaceae | Prevotellaceae UCG-004 | NA |
| ASV604 | 17.1 | -2.01 | 0.70 | 0.0970 | Firmicutes | UCG-010 | NA | NA |
| ASV321 | 22.4 | -1.95 | 0.53 | 0.0159 | Firmicutes | Oscillospiraceae | UCG-005 | NA |
| ASV225 | 43.3 | -1.85 | 0.49 | 0.0115 | Firmicutes | [Eubacterium] coprostanoligenes group | NA | NA |
| ASV177 | 44.0 | -1.40 | 0.48 | 0.0970 | Firmicutes | [Eubacterium] coprostanoligenes group | NA | NA |
| ASV57 | 121.7 | -1.34 | 0.46 | 0.0970 | Firmicutes | Oscillospiraceae | UCG-005 | NA |
| ASV77 | 157.9 | -1.07 | 0.35 | 0.0758 | Firmicutes | Lachnospiraceae | Shuttleworthia | NA |
| ASV44 | 145.9 | 0.78 | 0.24 | 0.0418 | Firmicutes | [Eubacterium] coprostanoligenes group | NA | NA |
| ASV288 | 39.6 | 0.91 | 0.30 | 0.0758 | Firmicutes | UCG-010 | NA | NA |
| ASV308 | 33.3 | 1.84 | 0.63 | 0.0970 | Firmicutes | Oscillospiraceae | UCG-005 | NA |
| ASV452 | 35.1 | 1.92 | 0.63 | 0.0758 | Firmicutes | Oscillospiraceae | NK4A214 group | NA |
| ASV205 | 63.1 | 2.34 | 0.61 | 0.0109 | Firmicutes | f__[Eubacterium] coprostanoligenes group | NA | NA |
| ASV2694 | 9.2 | 3.84 | 1.15 | 0.0418 | Proteobacteria | NA | NA | NA |
| ASV330 | 16.7 | 3.86 | 1.08 | 0.0194 | Euryarchaeota | Methanobacteriaceae | Methanobrevibacter | ruminantium |
| ASV16 | 167.8 | 4.56 | 1.14 | 0.0083 | Fusobacteriota | Leptotrichiaceae | NA | NA |
| ASV1255 | 21.9 | 4.91 | 1.34 | 0.0159 | Proteobacteria | NA | NA | NA |
| ASV12 | 28.0 | 5.36 | 1.33 | 0.0083 | Fusobacteriota | Fusobacteriaceae | Fusobacterium | necrophorum |
| ASV446 | 22.4 | 7.19 | 1.30 | 0.0000 | Proteobacteria | NA | NA | NA |

<sup>a</sup> Positive log<sub>2</sub> fold change values indicate that an ASV was more abundant in the Pregnant cows and vice versa.

#### Interaction of the network structure of the uterine microbiota between open and pregnant cows

After observing the community structure and compositional differences in the uterine microbiota between the two groups of cows, we next used ecological network modeling to analyze directed interactions among all observed genera. As shown in the network plots (Fig 4), cows that failed to become pregnant via AI had a noticeably distinct interaction network structure as compared to cows that became pregnant. Compared to the network structure observed in non-pregnant group, the complexity of the interaction network of the uterine microbiota from pregnant cows was reduced with a lower number of total genera that remained in the model. Despite having less intense interaction network structure, the total number of hubs connecting the interactions between genera was greater than the total number of hubs observed in the network structure of non-pregnant group (8 vs. 3 hubs). Overall, an equal proportion of positive and negative infractions between genera was detected in both network models.

**Figure 4.**
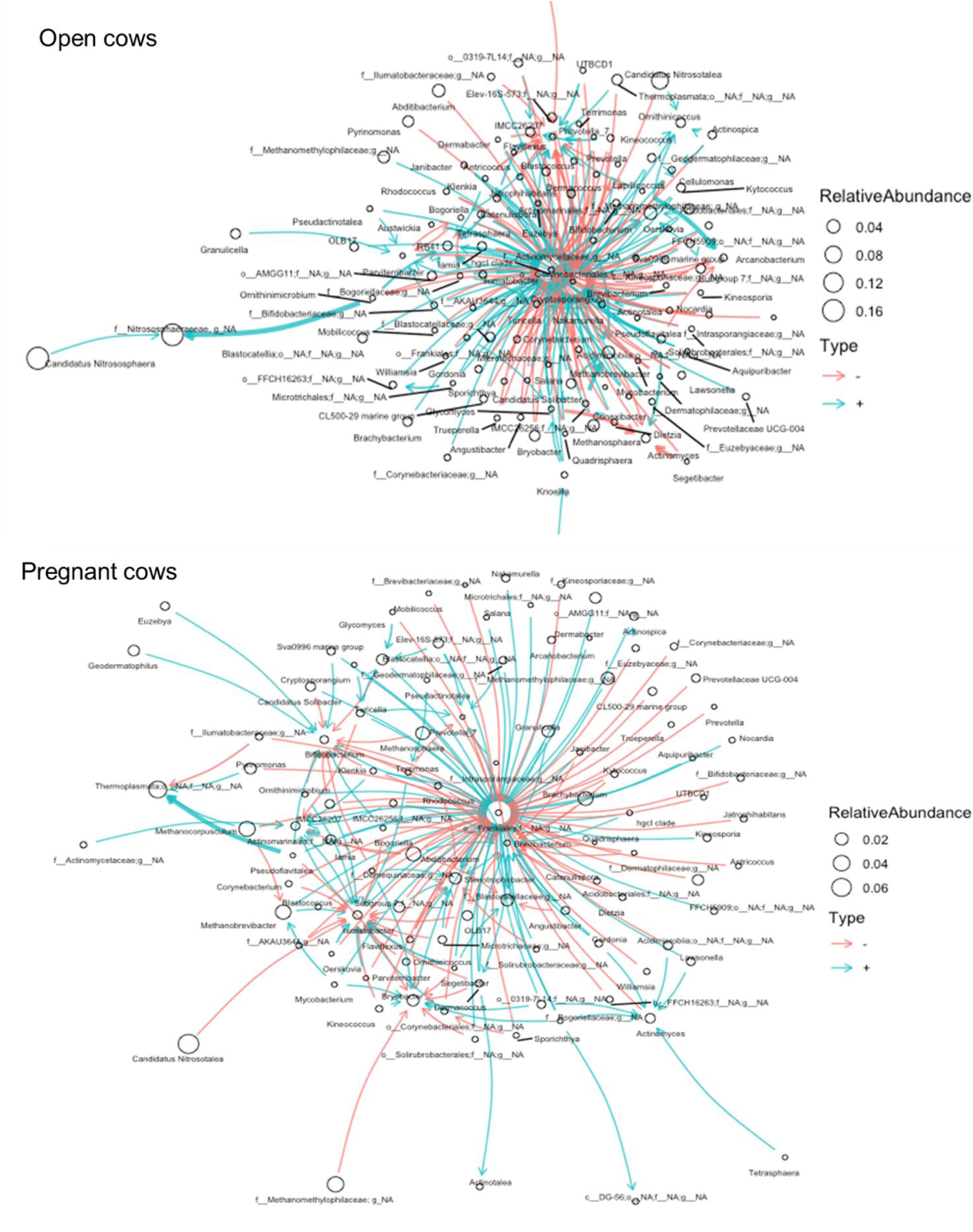
Ecological network of observed bacterial genera in uterine swab samples of non-pregnant (open, n = 31) and pregnant cows (n = 29) calves.

### Comparing the vaginal and uterine microbiota in cows

Given that vaginal and uterine swabs were collected simultaneously from the same animal, we compared the microbial composition and diversity between the vaginal and uterine microbiota. Overall, the microbial community in the uterus was distinct from that of the vagina in terms of community structure (R^2^ = 0.09; *P* < 0.0001) (Fig. 5A), richness, and diversity (*P* ≤ 0.04) (Fig. 5B). As expected, the diversity and richness of the vaginal microbiota were significantly greater than the uterine microbiota. Both reproductive organs were colonized by the same dominant bacteria phyla but the relative abundance of some of them (Bacteroidota, Actinobacteriota and Fusobacteriota) varied between the two sites (Fig. 5C). Despite these differences, a total of 6,481 ASVs (26% of overall ASVs) were shared between the vaginal and uterine microbiota (Fig. 5D). As shown in the heatmap (Fig. 6), the vast majority of the 100 most abundant ASVs were present in both vaginal and uterine swabs with similar frequency and abundance. Only a few taxa such as ASV8 (*Helicococcus ovis*), ASV79 (*Corynebacterum renale*) and ASV54 (*Corynebacterium*) were almost exclusively present in the uterus. The ASV43 ([Eubacterium] coprostanoligenes group) was the only taxon exclusively found in vaginal samples. Overall, a significant number of bacterial species are present in both the vagina and uterus of cattle.

**Figure 5.**
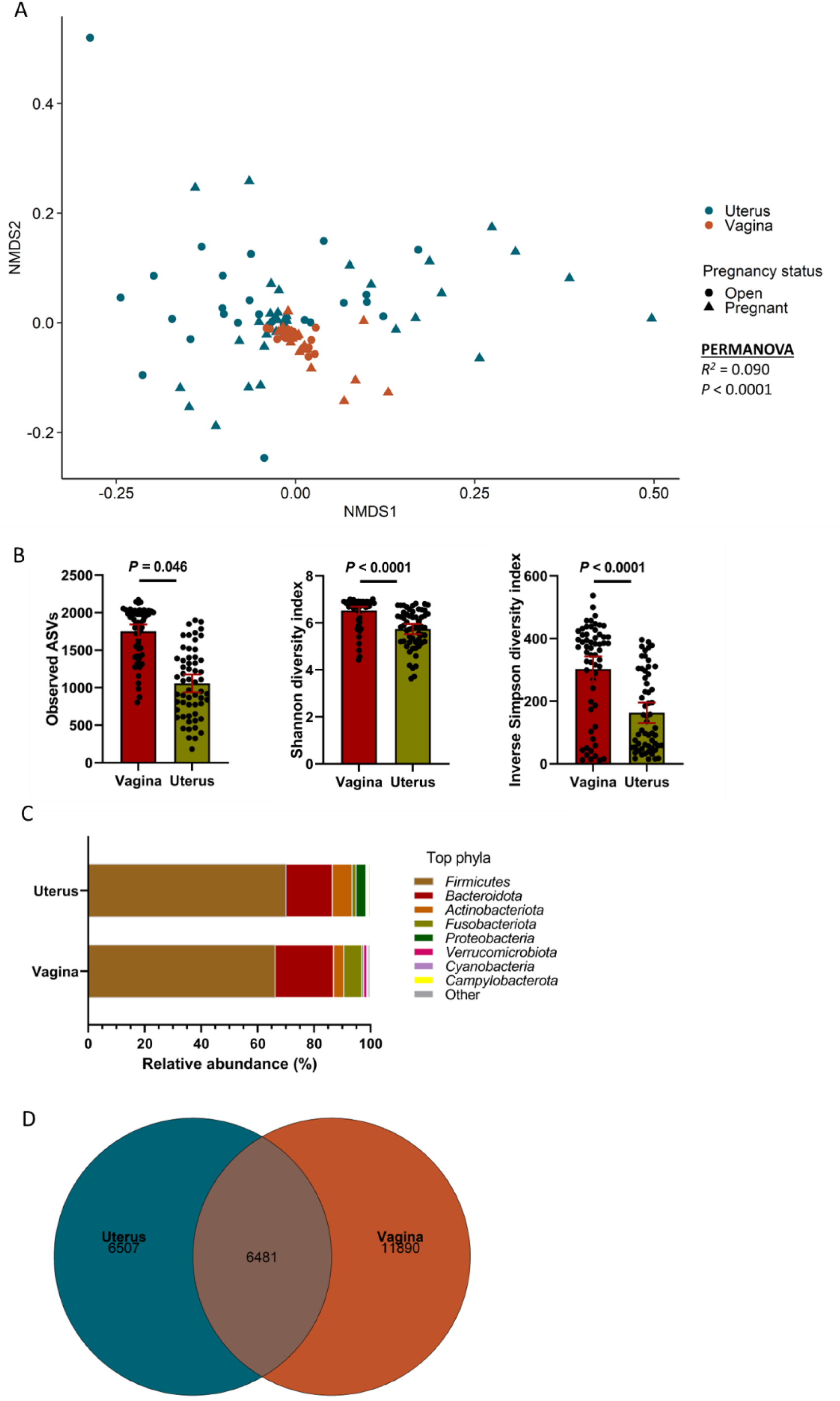
Beta and alpha diversity, and composition (phylum level) of the vaginal and uterine microbiota from cows (N = 60) that become pregnant or non-pregnant via AI. (**A**) Nonmetric multidimensional scaling (NMDS) plots of the Bray–Curtis dissimilarities, (**B**) number of observed amplicon sequence variants (ASVs), and Shannon and inverse Simpson’s diversity index, (**C**) Percent relative abundance of the eight most relatively abundant phyla, (**D**) | Venn diagram displaying the number of shared and unique ASVs between uterine and vaginal microbiota.

**Figure 6.**
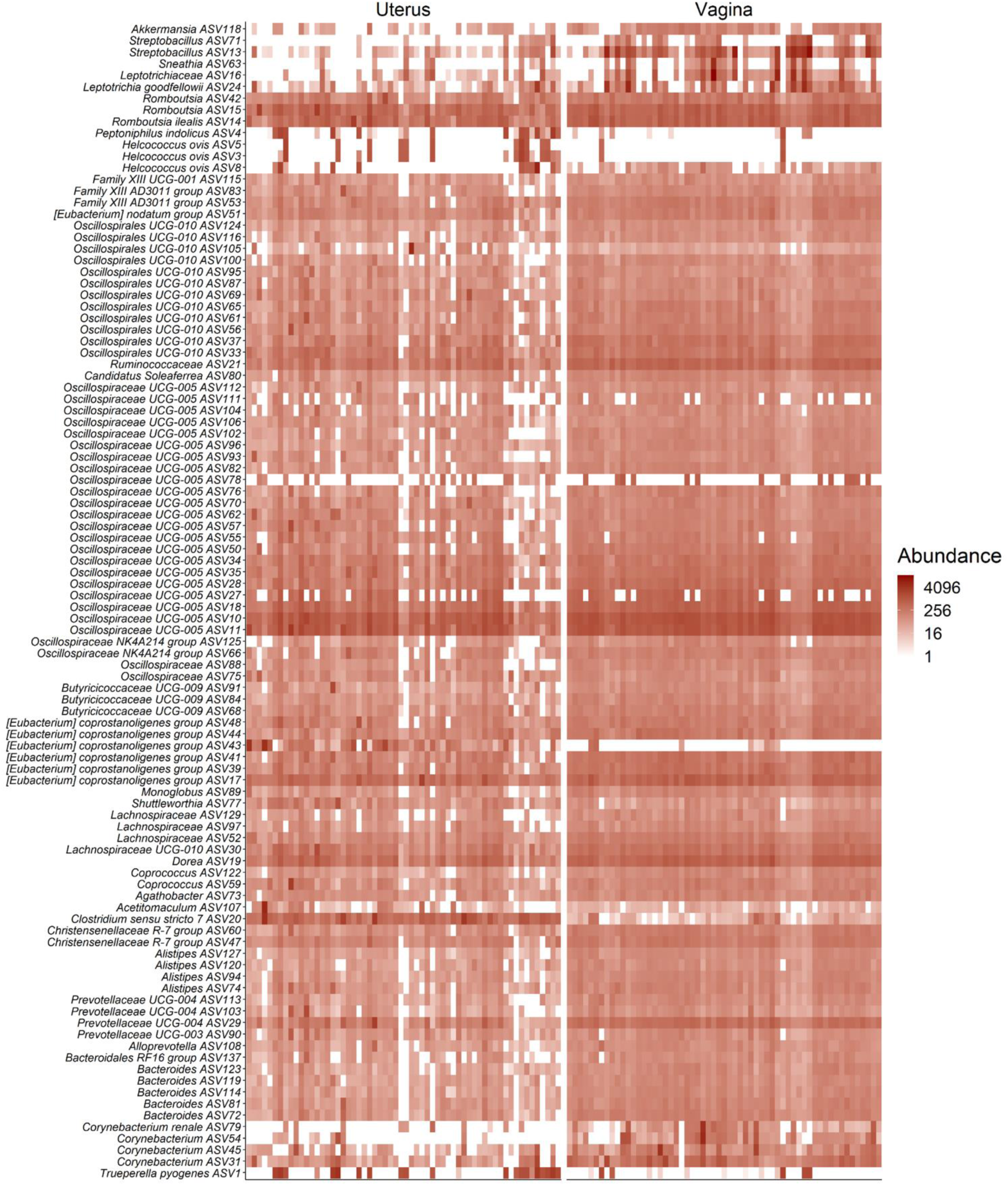
Heat map showing the 100 most abundant ASVs (log4) overall by uterine and vaginal swabs obtained from cows (N = 60) that become pregnant or non-pregnant via AI.

### Comparing the vaginal microbiota between heifers and cows

There was a significant and very large difference in the community structure of the vaginal microbiota between cows and heifers (R^2^ = 0.688; *P* < 0.001) (Fig. 7A). Vaginal microbial species richness and diversity were much greater in cows than heifers (Fig. 7B). Such differences were also reflected in the microbial composition, particularly the relative abundance of top bacterial phyla. The vaginas of cows were predominantly colonized by species within the phyla Firmicutes and Bacteroidota, whereas the vaginas of heifers were dominated equally by Firmicutes and Actinobacteriota.

**Figure 7.**
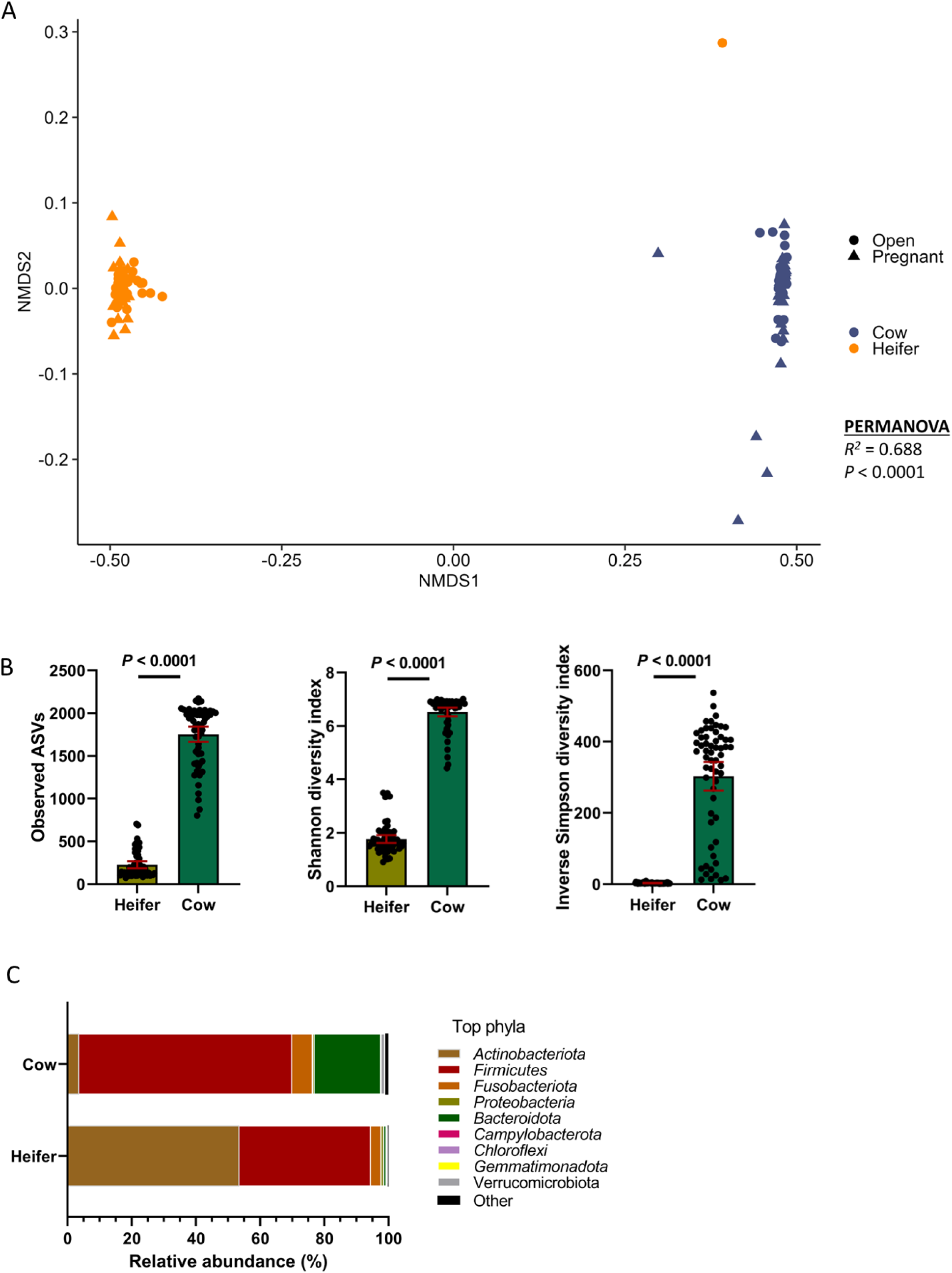
Beta and alpha diversity, and composition (phylum level) of the vaginal microbiota between heifers (N = 59) and cows (N = 60). (**A**) Nonmetric multidimensional scaling (NMDS) plots of the Bray–Curtis dissimilarities, (**B**) number of observed amplicon sequence variants (ASVs), and Shannon and inverse Simpson’s diversity index, (**C**) Percent relative abundance of the eight most relatively abundant phyla.

### Isolation and identification of vaginal and uterine bacterial isolates using aerobic and anaerobic culturing

#### Vaginal bacterial isolates

The vaginal swabs of 84 animals were plated aerobically and anaerobically which resulted in the recovery of 512 bacterial isolates, representing 52 genera within the phyla Firmicutes (58%), Proteobacteria (28%), Actinobacteria (12%), and Deferribacterota (2%) (Table 6). *Streptococcus* (37%)*, Bacillus* (19%)*, Escherichia* (7%)*, Staphylococcus* (5%), and *Corynebacterium* (4%) were the most frequently isolated genera. From the aerobic culturing of 64 vaginal swabs plated on MRS (semi-selective for lactic acid bacteria) and CB (non-selective) agar plates, 270 bacterial isolates (MRS = 92; CB = 178 isolates) were recovered (Table 6). These isolates were mainly from the three phyla Firmicutes (73%), Proteobacteria (16%), and Actinobacteria (11%), and 34 different genera with *Bacillus* (28%), *Streptococcus* (26%), *Escherichia* (9%), *Corynebacterium* (6%), *Trueperella* (4%) and *Staphylococcus* (4%) being the most abundant genera.

**Table 6.** Bacterial isolates from aerobic and anaerobic culturing of vaginal swab samples collected from yearling heifers and cows of various ages. Blue highlighted genera were only present in heifers and green highlighted genera were only found in cows.

| Phylum | Genus | Aerobic |  |  | Anaerobic |  |  |  | Total |
| --- | --- | --- | --- | --- | --- | --- | --- | --- | --- |
|  |  | CB | MRS | Subtotal | Blood | MRS | WC | Subtotal |  |
| Firmicutes | *Abyssicoccus* | ND | ND | ND | ND | ND | 1 | 1 | 1 |
| Firmicutes | *Aerococcus* | 3 | 1 | 4 | 2 | 3 | 5 | 10 | 14 |
| Firmicutes | *Alkalihalobacillus* | 5 | ND | 5 | ND | ND | ND | ND | 5 |
| Firmicutes | *Bacillus* | 39 | 37 | 76 | 8 | 1 | 13 | 22 | 98 |
| Firmicutes | *Brevibacillus* | 1 | ND | 1 | ND | ND | ND | ND | 1 |
| Firmicutes | *Clostridium* | ND | ND | ND | ND | ND | 1 | 1 | 1 |
| Firmicutes | *Enterococcus* | ND | ND | ND | 1 | 2 | 1 | 4 | 4 |
| Firmicutes | *Facklamia* | 2 | ND | 2 | 1 | 1 | 1 | 3 | 5 |
| Firmicutes | *Helcococcus* | 1 | ND | 1 | ND | ND | ND | ND | 1 |
| Firmicutes | *Heyndrickxia* | 1 | 4 | 5 | ND | ND | ND | ND | 5 |
| Firmicutes | *Hutsoniella* | 2 | ND | 2 | 1 | ND | 3 | 4 | 6 |
| Firmicutes | *Kurthia* | ND | ND | ND | ND | ND | 1 | 1 | 1 |
| Firmicutes | *Lacrimispora* | ND | ND | ND | ND | 1 | ND | 1 | 1 |
| Firmicutes | *Lactiplantibacillus* | ND | ND | ND | ND | 1 | ND | 1 | 1 |
| Firmicutes | *Lactobacillus* | ND | ND | ND | 1 | 1 | ND | 2 | 2 |
| Firmicutes | *Limosilactobacillus* | ND | ND | ND | ND | 1 | ND | 1 | 1 |
| Firmicutes | *Lysinibacillus* | 1 | 2 | 3 | ND | ND | ND | ND | 3 |
| Firmicutes | *Macrococcus* | 3 | ND | 3 | ND | ND | ND | ND | 3 |
| Firmicutes | *Neobacillus* | ND | ND | ND | 1 | ND | ND | 1 | 1 |
| Firmicutes | *Niallia* | 1 | 1 | 2 | 1 | ND | ND | 1 | 3 |
| Firmicutes | *Paenibacillus* | 8 | ND | 8 | 1 | ND | 3 | 4 | 12 |
| Firmicutes | *Peptoniphilus* | ND | ND | ND | 4 | 1 | 2 | 7 | 7 |
| Firmicutes | *Peribacillus* | ND | ND | ND | ND | ND | 1 | 1 | 1 |
| Firmicutes | *Periweissella* | ND | ND | ND | ND | ND | 1 | 1 | 1 |
| Firmicutes | *Priestia* | ND | 1 | 1 | ND | ND | ND | ND | 1 |
| Firmicutes | *Robertmurraya* | 1 | ND | 1 | ND | ND | ND | ND | 1 |
| Firmicutes | *Rossellomorea* | ND | 1 | 1 | ND | ND | ND | ND | 1 |
| Firmicutes | *Staphylococcus* | 9 | 2 | 11 | 6 | 2 | 7 | 15 | 26 |
| Firmicutes | *Streptococcus* | 40 | 30 | 70 | 50 | 42 | 27 | 119 | 189 |
| Firmicutes | *Weizmannia* | ND | ND | ND | ND | 2 | ND | 2 | 2 |
| Proteobacteria | *Acinetobacter* | 5 | ND | 5 | ND | ND | ND | ND | 5 |
| Proteobacteria | *Actinobacillus* | 2 | ND | 2 | 1 | ND | 4 | 5 | 7 |
| Proteobacteria | *Atlantibacter* | ND | ND | ND | ND | ND | 1 | 1 | 1 |
| Proteobacteria | *Campylobacter* | ND | ND | ND | ND | ND | 1 | 1 | 1 |
| Proteobacteria | *Escherichia* | 15 | 9 | 24 | 2 | 6 | 2 | 10 | 34 |
| Proteobacteria | *Histophilus* | 1 | ND | 1 | ND | ND | ND | ND | 1 |
| Proteobacteria | *Hydrogenophaga* | 1 | ND | 1 | ND | ND | ND | ND | 1 |
| Proteobacteria | *Kosakonia* | ND | 1 | 1 | ND | ND | ND | ND | 1 |
| Proteobacteria | *Lelliottia* | ND | ND | ND | ND | ND | 1 | 1 | 1 |
| Proteobacteria | *Mannheimia* | 3 | ND | 3 | ND | ND | 3 | 3 | 6 |
| Proteobacteria | *Moraxella* | 3 | ND | 3 | ND | ND | ND | ND | 3 |
| Proteobacteria | *Proteus* | 1 | ND | 1 | 5 | 1 | 3 | 9 | 10 |
| Proteobacteria | *Pseudescerichia* | ND | 1 | 1 | ND | ND | ND | ND | 1 |
| Proteobacteria | *Shigella* | 1 | ND | 1 | 1 | ND | ND | 1 | 2 |
| Proteobacteria | *Thioalkalivibrio* | ND | ND | ND | ND | ND | 1 | 1 | 1 |
| Actinobacteria | *Actinomadura* | ND | 1 | 1 | ND | ND | ND | ND | 1 |
| Actinobacteria | *Arthrobacter* | 2 | ND | 2 | ND | ND | ND | ND | 2 |
| Actinobacteria | *Corynebacterium* | 14 | 1 | 15 | 2 | ND | 5 | 7 | 22 |
| Actinobacteria | *Rothia* | 1 | ND | 1 | ND | ND | ND | ND | 1 |
| Actinobacteria | *Streptomyces* | 1 | ND | 1 | ND | ND | ND | ND | 1 |
| Actinobacteria | *Trueperella* | 11 | ND | 11 | ND | ND | ND | ND | 11 |
| Deferribacteres | *Denitrovibrio* | ND | ND | ND | ND | ND | 1 | 1 | 1 |
| Total number of isolates |  | 178 | 92 | 270 | 88 | 65 | 89 | 242 | 512 |

Thirty-four vaginal swabs were plated and cultured anaerobically on blood, MRS and WC agar plates resulting in the recovery of a total of 242 bacterial isolates (Table 6). The 88 isolates recovered from blood plates were taxonomically assigned to 17 genera from 3 phyla: Firmicutes (88%), Proteobacteria (10%), and Actinobacteria (2%). The isolates recovered from MRS plates (n = 65) were from 14 different genera within the Firmicutes (89%) and Proteobacteria (11%) phyla. The WC plates yielded slightly more diverse bacterial species (n = 89) comprised of 24 different genera which were from 4 different phyla: Firmicutes (75%), Proteobacteria (18%), Actinobacteria (6%), and Deferribacterota (1%). Overall, 32 different genera within 4 phyla: Firmicutes (83%), Proteobacteria (13%), Actinobacteria (3%), and Denitrovibrio (1%) were recovered from anaerobic culturing. The five most prevalent genera recovered were *Streptococcus* (49%)*, Bacillus* (9%)*, Staphylococcus* (6%)*, Aerococcus* (4%), and *Escherichia* (4%).

#### Uterine bacterial isolates

Overall, 221 bacterial isolates were recovered from the uterine swabs (n = 44) that were cultured aerobically and anaerobically (Table 7). These isolates were from 29 different genera within the Firmicutes (52%), Proteobacteria (34%), and Actinobacteria (14%) phyla. *Streptococcus* (48%)*, Bacillus* (15%)*, Escherichia* (6%)*, Corynebacterium* (5%), and *Facklamia* (3%) were the predominant genera.

**Table 7.**
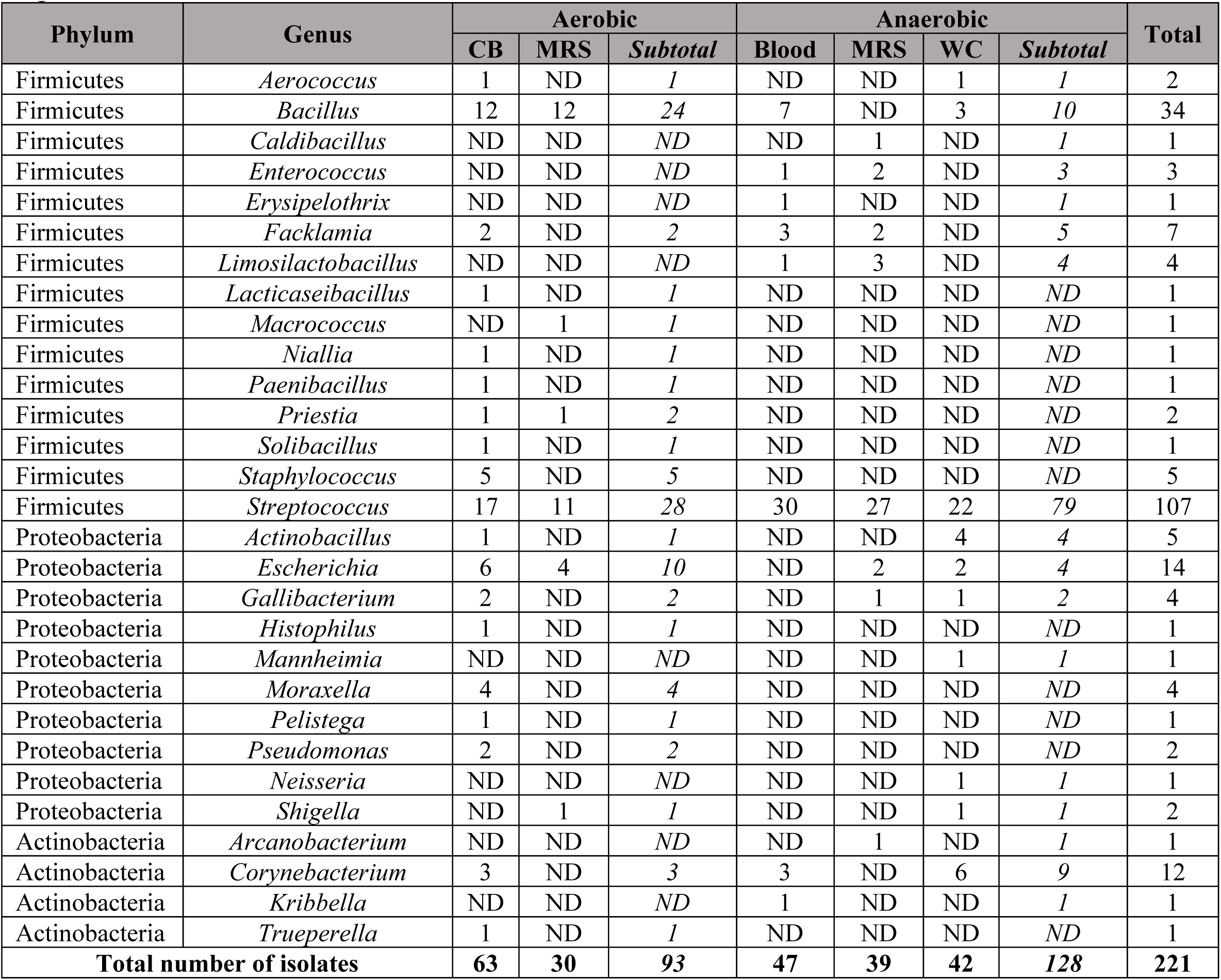
Bacterial isolates from aerobic and anaerobic culturing of uterine swab samples collected from cows of various ages.

Twenty-four uterine swabs were cultured aerobically on CB and MRS agar plates (Table 7) and yielded 93 bacterial isolates representing 21 genera. The five most predominant genera were *Streptococcus* (30%)*, Bacillus* (26%)*, Escherichia* (11%)*, Staphylococcus* (5%), and *Moraxella* (4%). The remaining 128 isolates were recovered from 34 uterine swabs after anerobic culturing on blood, MRS, and WC agar plates (Table 7). These isolates were from 17 bacterial genera within the Firmicutes (81%), Proteobacteria (10%), and Actinobacteria (9%) phyla. The most frequently isolated genera were *Streptococcus* (62%), *Bacillus* (8%), *Corynebacterium* (7%), *Facklamia* (4%), *Limosilactobacillus* (3%), *Actinobacillus* (3%), and *Escherichia* (3%).

### Antimicrobial susceptibility testing of selected vaginal and uterine isolates

The MICs of 41 different antibiotics from three different antibiotic panels were tested for 29 bacterial isolates (10 Gram-positive and 19 Gram-negative) from 6 different genera (Tables 8 and 9). Based on the CLSI breakpoints, most of the Gram-positive isolates were susceptible to all antibiotics tested, with *Staphylococcus epidermidis* (101.CB-6.VS) potentially showing resistance to erythromycin (Table 8). *Enterococcus hirae* 17168.V-An_MRS-A was resistant to amikacin, cephalothin, cefazolin, trimethoprim-sulfamethoxazole, clindamycin, and gentamicin. Among Gram-negative isolates tested, most *Escherichia coli* isolates exhibited intermediate resistance to cefazolin (Table 9), and one of them (18333.US_CB-C) was also resistant to doxycycline. The remaining Gram-negative isolates were susceptible to most of the antibiotics tested. Of note, several isolates such as *S. epidermidis* (101.CB-6.VS), *E. coli* (18333.US_CB-C), and *Histophilus somni* (20116.US_CB-A and 029V_CB-B), were not inhibited at the maximum concentration of antibiotics included in the panel, which makes it challenging to identify the resistance breakpoints. There are no available breakpoints for *Actinobacillus seminist* (306V_CB-B) for any of the 20 antibiotics tested. Five antibiotics that were tested on the Gram-positive panel (amikacin, marbofloxacin, pradofloxacin, enrofloxacin, and imipenem) and seventeen antibiotics from the Gram-negative panel (ceftiofur, clindamycin, danofloxacin, gamithromycin, erythromycin, clarithromycin, penicillin G, enrofloxacin, sulfadimethoxine, tiamulin, tildipirosin, tilmicosin, florfenicol, tulathromycin, tylosin tartrate, neomycin, and spectinomycin) had few to no breakpoint values documented for those isolates that we tested. Therefore, antibiotic resistance against these antibiotics could not be interpreted.

**Table 8.** MICs of antibiotics against Gram-positive bacterial isolates (n = 10) isolated from the uterus (n= 3) and vagina (n= 7) of beef cows and heifers^1^.

| Antibiotic | *Corynebacterium renale* (14121.US_CB-D) | *Staphylococcus chromogenes* (19197.US_CB-C) | *Staphylococcus chromogenes* (18207.US_CB-D) | *Corynebacterium renale* (14137.CB-1.VS) | *Enterococcus hirae* (17168.V-An_MRS-A) | *Staphylococcus aureus* (14177.V-An_MRS-A) | *Staphylococcus chromogenes* (247_CB-C.VS) | *Staphylococcus chromogenes* (240_CB-B.VS) | *Staphylococcus chromogenes* (306V_CB-A) | *Staphylococcus epidermidis* (101.CB-6.VS) |
| --- | --- | --- | --- | --- | --- | --- | --- | --- | --- | --- |
| Amikacin | ≤ 16 | ≤ 16 | ≤ 16 | ≤ 16 | > 32 | ≤ 16 | ≤ 16 | ≤ 16 | ≤ 16 | ≤ 16 |
| Ampicillin | ≤ 0.25 | ≤ 0.25 | ≤ 0.25 | ≤ 0.25 | = 2 | ≤ 0.25 | ≤ 0.25 | ≤ 0.25 | ≤ 0.25 | ≤ 0.25 |
| Amoxicillin-clavulanate | ≤ 0.25 | ≤ 0.25 | ≤ 0.25 | ≤ 0.25 | = 1 | ≤ 0.25 | ≤ 0.25 | ≤ 0.25 | ≤ 0.25 | ≤ 0.25 |
| Cephalothin | ≤ 2 | ≤ 2 | ≤ 2 | ≤ 2 | > 4 | ≤ 2 | ≤ 2 | ≤ 2 | ≤ 2 | ≤ 2 |
| Clindamycin | ≤ 0.5 | ≤ 0.5 | ≤ 0.5 | ≤ 0.5 | > 4 | ≤ 0.5 | ≤ 0.5 | ≤ 0.5 | ≤ 0.5 | ≤ 0.5 |
| Cefazolin | ≤ 2 | ≤ 2 | ≤ 2 | ≤ 2 | > 4 | ≤ 2 | ≤ 2 | ≤ 2 | ≤ 2 | ≤ 2 |
| Gentamicin | ≤ 4 | ≤ 4 | ≤ 4 | ≤ 4 | = 8 | ≤ 4 | ≤ 4 | ≤ 4 | ≤ 4 | ≤ 4 |
| Trimethoprim-sulfamethoxazole | ≤ 2 | ≤ 2 | ≤ 2 | ≤ 2 | ≤ 2 | ≤ 2 | ≤ 2 | ≤ 2 | ≤ 2 | ≤ 2 |
| Minocycline | ≤ 0.5 | ≤ 0.5 | = 1 | ≤ 0.5 | ≤ 0.5 | ≤ 0.5 | ≤ 0.5 | = 1 | = 1 | ≤ 0.5 |
| Chloramphenicol | ≤ 8 | ≤ 8 | ≤ 8 | ≤ 8 | ≤ 8 | ≤ 8 | ≤ 8 | ≤ 8 | ≤ 8 | ≤ 8 |
| Marbofloxacin | = 2 | ≤ 1 | ≤ 1 | = 2 | ≤ 1 | ≤ 1 | ≤ 1 | ≤ 1 | ≤ 1 | ≤ 1 |
| Erythromycin | ≤ 0.25 | = 0.5 | = 0.5 | ≤ 0.25 | ≤ 0.25 | ≤ 0.25 | ≤ 0.25 | = 0.5 | = 0.5 | > 4 |
| Pradofloxacin | ≤ 0.25 | ≤ 0.25 | ≤ 0.25 | ≤ 0.25 | ≤ 0.25 | ≤ 0.25 | ≤ 0.25 | ≤ 0.25 | ≤ 0.25 | ≤ 0.25 |
| Penicillin G | ≤ 0.06 | ≤ 0.06 | ≤ 0.06 | ≤ 0.06 | = 4 | ≤ 0.06 | ≤ 0.06 | ≤ 0.06 | ≤ 0.06 | ≤ 0.06 |
| Enrofloxacin | = 1 | ≤ 0.25 | ≤ 0.25 | = 0.5 | = 1 | ≤ 0.25 | ≤ 0.25 | ≤ 0.25 | ≤ 0.25 | ≤ 0.25 |
| Nitrofurantoin | ≤ 16 | ≤ 16 | ≤ 16 | ≤ 16 | = 32 | ≤ 16 | ≤ 16 | ≤ 16 | ≤ 16 | ≤ 16 |
| Imipenem | ≤ 1 | ≤ 1 | ≤ 1 | ≤ 1 | = 4 | ≤ 1 | ≤ 1 | ≤ 1 | ≤ 1 | ≤ 1 |
| Doxycycline | ≤ 0.125 | ≤ 0.125 | ≤ 0.125 | ≤ 0.125 | = 0.25 | ≤ 0.125 | ≤ 0.125 | > 0.5 | > 0.5 | ≤ 0.125 |
| Cefpodoxime | ≤ 2 | ≤ 2 | ≤ 2 | ≤ 2 | > 8 | ≤ 2 | ≤ 2 | ≤ 2 | ≤ 2 | ≤ 2 |
| Tetracycline | ≤ 0.25 | = 0.5 | ≤ 0.25 | ≤ 0.25 | = 0.5 | ≤ 0.25 | ≤ 0.25 | > 1 | > 1 | ≤ 0.25 |
| Cefovecin | = 0.25 | = 0.125 | = 0.125 | = 0.25 | > 8 | = 1 | = 1 | = 1 | = 1 | = 0.125 |
| Vancomycin | ≤ 1 | ≤ 1 | ≤ 1 | ≤ 1 | ≤ 1 | ≤ 1 | ≤ 1 | ≤ 1 | ≤ 1 | ≤ 1 |
| Oxacillin | = 0.5 | ≤ 0.25 | ≤ 0.25 | = 0.5 | > 2 | ≤ 0.25 | ≤ 0.25 | ≤ 0.25 | ≤ 0.25 | ≤ 0.25 |
| Rifampin | ≤ 1 | ≤ 1 | ≤ 1 | ≤ 1 | ≤ 1 | ≤ 1 | ≤ 1 | ≤ 1 | ≤ 1 | ≤ 1 |

<sup>1</sup>MIC values (µg/ml) presented in the table were obtained from the Sensititre Gram Positive Companion panel (COMPGP1F).
GREEN= no break points (isolate)
BLUE= no/few break points (antibiotic)
RED= resistant
DARK RED/WHITE TEXT= intrinsically resistant
YELLOW= highest test concentrations were lower than breakpoints
ORANGE= intermediate

**Table 9.**
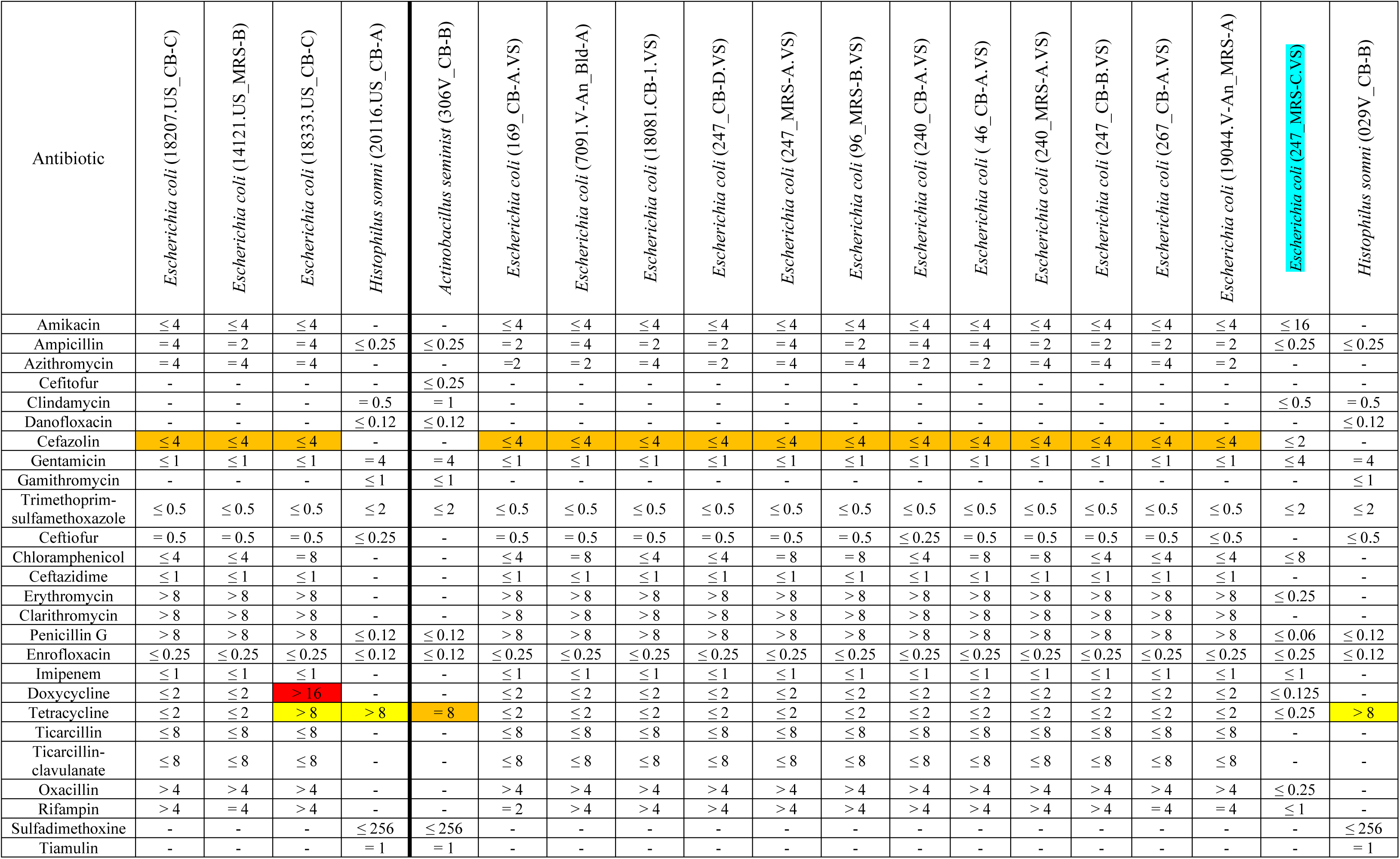
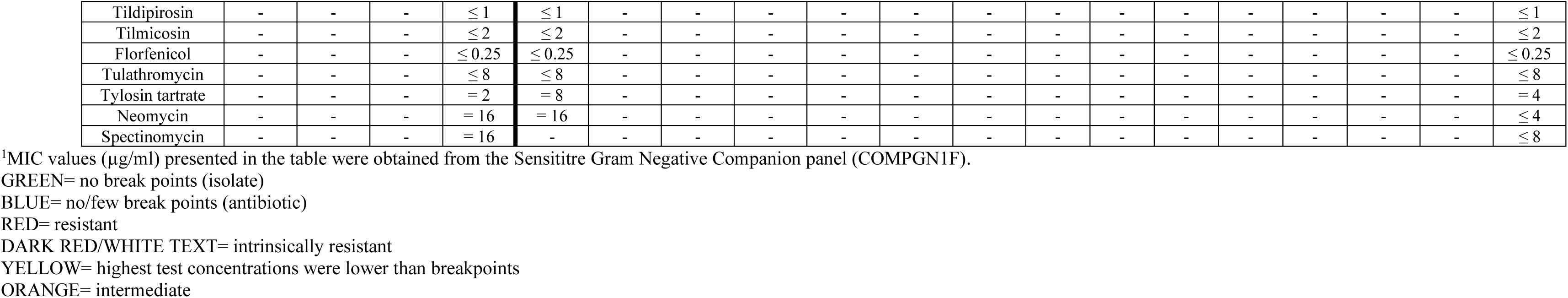
MICs of antibiotics against Gram-negative bacterial isolates (n = 19) isolated from the uterus (n= 4) and vagina (n= 15) of beef cows and heifers^1^.

## Discussion

Although there have been significant advances in AI, genetic selection, and feeding and management practices in the past several decades, reproductive failure persists as a significant problem affecting the beef and dairy cattle industries (Ayalon, 1978; Sheldon and Dobson, 2003; Reese et al., 2020). Increasing evidence indicates that the microbial communities residing within the female reproductive tract may play a critical role in fertility, and thus could be targeted for reducing reproductive failure in cattle. In this study, we evaluated the vaginal and uterine microbiota of beef heifers and cows that became pregnant via AI and those that did not to identify microbial signatures and differentially abundant taxa associated with pregnancy at the time of AI. We performed comparisons between the vaginal and uterine microbiota within cows, and between heifer and cow vaginal microbial communities. We also assessed the culturable fraction of the vaginal and uterine microbiota using extensive culturing (both anaerobic and aerobic) and isolated a relatively large number of bacterial strains with the intention of developing a future bacterial consortium that could be administered prior to or at the time of AI to enhance pregnancy outcomes. Additionally, we performed AST on selected vaginal and uterine bacterial isolates to gain insights into AMR present in the commensal bacteria residing within the female reproductive tract of cattle.

Overall, 16S rRNA gene sequencing results revealed that the vaginal microbiota at the time of AI remained the same in terms of community structure (heifers and cows), but its microbial richness and diversity (cows) tended to be reduced in cattle who become pregnant via AI compared to those that did not. Vaginal microbiota with a reduced richness and diversity have been considered “normal and healthy” and positively associated with reproductive health, pregnancy outcome, and lower preterm birth risk in women (Leitich et al., 2003; Aagaard et al., 2012; Freitas et al., 2017; Freitas et al., 2018). Thus, the lower microbial richness and diversity observed in the cow vaginal microbiota at the time of AI could be associated with pregnancy rate. However, no difference was observed in any of the alpha diversity metrics of the vaginal microbiota in virgin heifers which had not previously been bred at the time of sampling. This might be partially due to the fact that the virgin heifers harbor a vaginal community with significantly lower microbial richness and diversity compared to pregnant heifers (Amat et al., 2021) and cows (Fig 6B), and which would necessitates to use a greater number of animals in a study to detect differences. Thus, the association between the reduced microbial richness and diversity in yearling heifers and pregnancy rate warrants future study.

Moderate changes in the microbial composition at genus (cows) and ASV (heifers) levels were observed in the vagina between pregnant and non-pregnant groups. Cows that did not become pregnant had a greater relative abundance of *Monoglobus* and *Prevotellaceae* UCG-003 . *Monoglobus* spp., which belongs to the family *Ruminococcaceae,* is involved in degradation of roughage in the rumen (Pu et al., 2022) and have previously been reported to be among the most abundant genera in the vaginal microbiota of virgin yearling and pregnant beef heifers (Amat et al., 2021). There are no reports regarding the association of the *Monoglobus* in the vagina with reproductive health and pregnancy in either women or female cattle. Members of the *Prevotellaceae* family have been found to be associated with abnormal vaginal microbiota (Freitas et al., 2018) and reduced pregnancy rates in women (Romero et al., 2014). Thus, the role of vaginal *Monoglobus* and *Prevotellaceae* UCG-003 in bovine reproductive health and pregnancy outcomes will require further investigation. The 11 ASVs that were inversely associated with AI pregnancy rates in heifers, and which were only classified at the genus (7 out of 11), or higher taxonomic levels were from only three families: *Lachnospiraceae*, *Oscillospiraceae* and *Butyricicoccaceae* (Table 2). The abundance of vaginal *Lachnospiraceae* was reduced in antibiotic allergic pregnant women as compared to non-allergic pregnant women (Li et al., 2020). Both *Lachnospiraceae* and *Oscillospiraceae* were enriched in the vaginal microbiota of women with adenomyosis (Kunaseth et al., 2022), a condition where the endometrial tissue that forms the lining of the uterus grows into the muscle of the uterine wall and enlarges the uterus (Chernofsky, 2023). These reports and our results together suggest that a bovine vaginal microbiota with increased proportion of *Lachnospiraceae* and *Oscillospiraceae* at the time of breeding could be associated with reduced conception. The *Butyricicoccaceae* family encompasses butyric acid-producing species in the gut (Nava and Stappenbeck, 2011), and butyric acid produced by bacteria has been shown to regulate animal reproduction by directly regulating progesterone and 17β-estradiol secretion (Lu et al., 2017). The elevated relative abundance of vaginal *Butyricicoccaceae UCG-009* in non-pregnant heifers indicates that butyric acid-producing bacterial species in the reproductive tract may have role in influencing bovine reproduction. Taken together, our results highlight that the abundance of certain bacterial species in the vaginal tract of female cattle before or at the time of breeding may influence pregnancy outcomes.

An acceptable uterine swabbing technique for virgin yearling heifers was not available at the time of sampling, therefore, our uterine data was limited to only cows. Compared to the vaginal microbiota, more noticeable differences were observed in the uterine microbiota between pregnant and non-pregnant cows. These differences include a distinct community structure and interaction network structure, and differentially abundant taxa between the two groups. Both pregnant and non-pregnant cows had similar alpha diversity metric values at the time of AI. Cows that became pregnant harbored noticeably different interaction networks among the observed genera from the non-pregnant cows. Although it is challenging to make inferences on the biological meaning of ecological network modeling, active interactions with balanced positive (cooperation) and negative (competition) interconnectivity between different microbial species are important for maintaining the stability and functional features of the gut (Faust et al., 2012; Foster et al., 2017; Venturelli et al., 2018) and respiratory microbiota (Amat et al., 2023). Distinctive microbial relative co-abundance networks were observed in the vaginal microbiota between penicillin allergic and non-allergic women (Li et al., 2020). Accordingly, our network results suggest that directionality of the interactions and cooperative and competitive interconnectivity between different uterine microbial species may have implications in female fertility and pregnancy.

Among the 28 differentially abundant ASVs in the uterine microbiota, only three were assigned to a species. Two of these were *Methanobrevibacter ruminantium* (ASV330) and *Fusobacterium necrophorum* (ASV12) both of which were enriched in pregnant cows. Although we have previously reported that the methanogenic archaeal genus *Methanobrevibacter* is among the top 40 most relatively abundant genera present in the vaginal microbiota of pregnant heifers (Amat et al., 2021), and that *M. ruminantium* is present in the vagina and uterus of beef heifers (Winders et al., 2023), it is interesting to observe that this species has a positive correlation with pregnancy outcome in cows. The higher relative abundance of *F. necrophorum* observed in the uterus of cows that become pregnant is surprising as this species is commonly known as an opportunistic pathogen associated with several bovine diseases including liver abscesses and foot root, both of which have a negative effect on cattle performance and profitability in the beef cattle industry (Nagaraja and Chengappa, 1998; Nagaraja et al., 2005). *F. necrophorum* has also been associated with abortion in cattle (Clothier and Anderson, 2016; Reichel et al., 2018). However, new evidence indicates that *F. necrophorum* in the bovine urogenital tract may not always be harmful but could be a commensal bacterial species with a positive role in reproductive health and fertility. For example, *F. necrophorum* was isolated in the seminal microbiota of healthy beef bulls (Webb et al., 2023). Recently we identified *Fusobacterium* was one of the most relatively abundant genera (0.6% relative abundance) in the uterus of healthy beef heifers (Winders et al., 2023) and in the semen (26%) of healthy bulls (Webb et al., 2023). In the present study, the uterus of pregnant cows harbored 125-fold greater relative abundance of *Fusobacterium* as compared to non-pregnant cows at the time of AI (0.63% vs. 0.005%). According to a recent comparative genomic analysis of different *F. necrophorum* strains (Bista et al., 2022) and the site-specific genome adaptations reported in *Lactobacillus* spp. (Pan et al., 2020), it is likely that *F. necrophorum* species that colonize the reproductive tract do not contain the virulence genes found in the pathogenic strains implicated in the liver-abscesses, foot rot, or abortion. Thus, understanding the role of *Fusobacterium necrophorum* in cattle reproductive health and fertility, a genomic comparison between the *F.necrophorum* isolates isolated from healthy cow uterus and from clinical infections should be the focus of future studies.

Two *Oscillospiraceae* ASVs (UCG-005 ASV303 and NK4A214 ASV308) representing uncultured genera were also more abundant in the pregnant cow uterus. However, a different ASV classified as *Oscillospiraceae* UCG-005 was enriched in the vagina of non-pregnant heifers, and four ASVs within the family *Oscillospiraceae* (ASV592, 401, 321 and 57) were also enriched in the uterus of non-pregnant cows. This suggests that the family *Oscillospiraceae* could encompass multiple species or strains found in the reproductive tract of female cattle that may have anti-pregnancy properties. Similar to the observation in the cow vagina, the relative abundance of some taxa within the genera *Monoglobus* (ASV644), *Prevotella-7* (ASV385), and *Prevotellaceae* UCG-004 (ASV103) were significantly elevated in non-pregnant cow uterine samples, suggesting that *Monoglobus* and *Prevotella* spp. may negatively affect cattle fertility. ASV957, which was identified as *Faecalibacterium prausnitzii,* was more abundant in the uterus of non-pregnant cows. This bacterium consumes acetate and produces butyrate and other bioactive anti-inflammatory modulators (Sokol et al., 2008). A higher proportion of *F. prausnitzii* taxa in the uterus of non-pregnant cows and the enrichment of vaginal *Butyricicoccaceae* UCG-009 in non-pregnant heifers suggest that butyrate-producing bacteria in the urogenital tract of female cattle may have adverse effects on the reproductive health and pregnancy outcome. Non-pregnant cows also harbored a greater relative abundance of ASVs identified as *Rikenellaceae* RC9 gut group (ASV506 and *Christensenellaceae* R-7 group ASV6112 in their uterus. Although both of these genera are part of the common microbiota present in the cattle gut (Winders et al., 2023), and *Rikenellaceae* RC9 gut group also colonizes the bovine uterus (Yagisawa et al., 2023), there are no reports regarding their association with reproductive infections or fertility in cattle. Overall, our results, for the first time, show the presence of pregnancy associated microbial taxa among the bovine uterine microbiota.

Overall, the vaginal microbial community structure, species richness, diversity, and composition were drastically different between heifers and cows. Cows harbored a significantly richer and more diverse vaginal microbiota dominated by species within the phylum Firmicutes as compared to heifers (Fig. 7). In our recent study, we characterized the vaginal microbiota associated with 30-h old calves born from dams who were originated from the same farm as the cows used in the present study (Luecke et al., 2023). These newborn calves harbored vaginal microbiota with a lower richness (Total mean observed ASVs < 350) and diversity (Shannon diversity index < 3) compared to vaginal microbiota observed in heifers and cows in the present study. These results indicate that cattle age, reproductive developmental stage, sexual activity, and number of births have significant influence on the vaginal microbiome (Moreno et al., 2017; Mirzaei et al., 2023). Apart from these factors, the housing environment and feed could be contributing to the different vaginal microbiota observed between heifers and cows as these two cohorts were housed in two different environments; cows were on the pasture while heifers were house in an enclosed animal facility.

As expected, the structure, richness and diversity of the vaginal and uterine microbiota were significantly different from each other with the uterus having a less rich and diverse as compared to vagina. This might be due to physiological, immunological, microbiological, and anatomical differences in the vagina and uterus (Nguyen et al., 2014; Agostinis et al., 2019; Amat et al., 2022). Despite these differences, 26% of ASVs were shared between the vaginal and uterine microbiota, and the vast majority of the 100 most abundant ASVs were present in both sites. Such similarities between the vaginal and uterine microbiota suggest that bacterial species at the mucosal surface of the vagina may ascend into the uterus despite the difference in oxygen availability and other physiochemical properties between the two anatomical sites. Thus, future studies aimed at developing probiotics to enhance fertility and to be delivered *in utero* should consider those beneficial bacterial species present in the vaginal tract as well.

Considering that 89% and 57% of the differentially abundant taxa in the uterine samples were not classified at the genus and species levels, respectively, and that a similar proportion of vaginal bacteria were not classified at the genus level in both heifers and cows, we also used a comprehensive culturing of vaginal and uterine samples. We isolated and identified a total of 733 bacterial isolates (512 vaginal and 221 uterine isolates) using four different agar media under anaerobic and aerobic culturing conditions. Isolates from 52 different bacterial genera within 4 different phyla were recovered from aerobically and anaerobically (270 vs. 242) from vaginal swabs. The uterine samples yielded fewer isolates aerobically and anaerobically (93 vs. 128) and from fewer unique genera (29). Overall, the culturable fraction of the vaginal and uterine microbiota at both phylum and genus levels was noticeably different from that identified by 16S rRNA gene sequencing. Firmicutes (58%), Proteobacteria (28%) and Actinobacteria (12%) were the most frequently identified phyla among the cultured bacterial isolates. However, 16S rRNA gene sequencing showed that almost 95% vaginal microbiota was composed of Actinobacteria and Firmicutes in almost equal proportions. None of the prevalent genera (*Streptococcus*, *Bacillus*, *Escherichia,* and *Staphylococcus*) identified by culturing of vaginal swabs were among the 20 most relatively abundant genera in the vagina microbiota based on 16S rRNA gene sequencing. The most abundant genera in the uterine microbiota were also vastly different between culturing and sequencing. For example, we did not recover any species within *Leptotrichiaceae*, *Oscillospiraceae,* or *Lachnospiraceae* families or *Fusobacterium* and *Prevotella* genera by culturing (Table S1). Thus, results highlight a need for culturing with specialized growth media that support a wider range of aerobic and anaerobic species, and under conditions that can better replicate the physiological and mucosal properties of the bovine reproductive tract. One of the factors that may have limited the recovery of some bacterial species from the vaginal and uterine swabs was the freezing of the samples prior to culturing. It has been reported that freezer storage can influence viable bacterial cell recovery, and that different species may tolerate freezing and the cryoprotectant differently (Bircher et al., 2018; Webb et al., 2023).

To the best of our knowledge, this is the first study to characterized culturable portion of the female bovine reproductive microbiota. One of the interesting and important finding from our culture results is that both vaginal and uterine microbial communities in healthy female cattle harbor bacterial species associated with diseases. These bacteria species include *H. somni* (bovine respiratory disease) (Griffin et al., 2010), *Moraxella bovis* (pinkeye) (Loy et al., 2021), *Moraxella bovoculi* (pinkeye) (Loy et al., 2021), *Staphylococcus aureus* (mastitis) (Campos et al., 2022), and *Trueperella abortisuis* (abortion) (Alssahen et al., 2020) (Supplemental Table 1). Other genera such as *Shigella*, *Streptococcus*, and *Mannheimia* that contain pathogenic species were also identified. The presence of these potentially pathogenic bacteria within the vaginal and uterine microbial community begs the question of whether these bacteria are transferred to the offspring during birth or to the bulls during natural breeding (Luecke et al., 2022).

Antimicrobial resistance has become a pressing issue in both animal and public health, particularly with regard to dissemination through bovine-associated pathogenic and commensal bacteria (Cameron and McAllister, 2016). Therefore, surveillance of the resistome (the collection of AMR genes associated with the microbial community of a particular environment) is important to mitigate the emergence and spread of AMR (Ma et al., 2021). The microbial continuum along the female reproductive tract is an important source of seeding for the offspring calf (Amat et al., 2022; Luecke et al., 2022; Poole et al., 2023) and bull reproductive tract microbiota (Luecke et al., 2022). Considering that bacterial may be transfered from the female reproductive tract to the bull, then to other females, and ultimately to their offsprings (Luecke et al., 2022; Luecke et al., 2023), it is critical to investigate AMR in commensal bacteria present in the female reproductive tract of cattle. Our AST results for 29 aerobic vaginal and uterine isolates revealed that only one *Escherichia coli* isolate showed resistance, and majority of remaining isolates were susceptible to the antibiotics tested, suggesting that the commensal bacteria within the female urogenital tract harbor fewer antibiotic resistant bacteria than anticipated. However, given that there were no breakpoints available for 21 antibiotics (5 antibiotics in the Gram-positive panels and 16 antibiotics in Gram-negative panels), our AST results need to be interpreted cautiously. A larger set of bacterial isolates with bovine specific antibiotic panels, and if possible, shotgun metagenomic sequencing would be needed to fully determine the scope of the resistome associated with the female reproductive microbiome.

## Conclusions

Although the structure, species richness and diversity of the vaginal microbiota did not differ in heifers that failed to become pregnant via AI compared to heifers that did, certain taxa were found to be associated with pregnancy. Although no larger differences in the microbial community structure were noted, the vagina microbiota of non-pregnant cows tended to have greater richness and diversity compared to pregnant cows.

Twenty-eight taxa were differentially abundant in the uterine microbiota between the two groups of cows, including *Methanobrevibacter ruminantium* and *F. necrophorum* which were positively correlated with pregnancy. Our culturing results revealed that a relatively diverse assortment of bacterial species, mostly comprised of aerobic and anaerobic Gram-positive bacteria, is present in the bovine female reproductive tract. Most of the vaginal and uterine bacterial isolates screened for antimicrobial resistance did not show resistance. Our identification of pregnancy-associated vaginal and uterine microbial taxa at the time of breeding highlights the possibility of developing reproductive microbiome-targeted strategies to enhance fertility in beef cattle.

## Abbreviations

AI: artificial insemination
AMR: antimicrobial resistance
AST: antimicrobial susceptibility test(ing)
ASV: amplicon sequencing variant
BHI: brain heart infusion media
BHIg: brain heart infusion media with 20% glycerol
BLAST: basic local alignment search tool
bp: base pair
CB: Columbia blood agar
CIDR: controlled internal drug release
CLSI: Clinical and Laboratory Standards Institute
EDTA: ethylenediaminetetraacetic acid
IACUC: Institutional Animal Care and Use Committee
max.: maximum
MIC: minimum inhibitory concentration
min.: minimum
MRS: De Man, Rogosa, and Sharpe media
MRSg: De Man, Rogosa, and Sharpe media with 20% glycerol
PBS: phosphate buffered saline
PERMANOVA: permutational multivariate analysis of variance
qPCR: quantitative polymerase chain reaction
RDP classifier: ribosomal database project
SD: standard deviation
SSU: small subunit
TCEP: Tris (2-carboxyethyl) phosphine
TE: tris ethylenediaminetetraacetic acid
WC: Wilkins Chalgren media

## Acknowledgements

We thank Kelli Maddock at the North Dakota State University Veterinary Diagnostic Laboratory for her assistance in antimicrobial susceptibility testing. We acknowledge Godson Aryee for his assistance with the ecological network analysis. We also thank Layla King, Yssi Entzie and Jessica Syring in the Department of Animal Science, and staff at Animal Nutrition and Physiology Center (ANPC), NDSU for their assistance with animal care. We thank the animal caring staff at the Central Grassland Research and Extension Center (CGREC), NDSU.

## Author contributions

S.A. and C.R.D., Conceiving the idea, designing the study, and providing supervision; S.A., A.K.W, C.R.D. K.K.S, Cattle management; S.A., C.R.D., K.N.S., J.L.H., K.A.B., Animal care and sample collections; E.M.W., K.N.S., B.P and S.A., Sample processing; D.B.H., S.A., E.M.W., Bioinformatics analysis and data and statistical analysis; E.M.W., and S.A., Manuscript writing; S.A., D.B.H., E.M.W., K.N.S., A.K.W., and C.R.D., Manuscript review and editing. All authors have read and agreed to the published version of the manuscript.

## Conflict of interest

The authors declare that the research conducted in this study was absent of any conflict of interest.

## Funding

The work presented in this study was financially supported by the North Dakota Agricultural Experiment Station as part of a start-up package for SA, as well as a 2021-2022 North Dakota State University EPSCoR STEM Research and Education Funding-Seed Award.

## Data availability

Raw sequence data are available from the NCBI Sequence Read Archive under BioProject accession PRJNA976303. All other data supporting the findings of this study are presented within the paper.

---

**Supplemental Table S1.** A list of all bacterial isolates recovered, their growth media and oxygen condition, and their anatomic origin.

| Phylum | Bacteria Name | Number of Isolates |  | Isolate Origin |  | Media |  |  |
| --- | --- | --- | --- | --- | --- | --- | --- | --- |
|  |  | Aerobic | Anaerobic | Uterus | Vagina | CB/Blood | MRS | WC |
| Firmicutes | *Abyssicoccus albus* | - | 1 | - | 1 | - | - | 1 |
| Proteobacteria | *Acinetobacter lwoffii* | 3 | - | - | 3 | 3 | - | - |
| Proteobacteria | *Acinetobacter seohaensis* | 2 | - | - | 2 | 2 | - | - |
| Proteobacteria | *Actinobacillus seminis* | 3 | 9 | 5 | 7 | 4 | - | 8 |
| Actinobacteria | *Actinomadura formosensis* | 1 | - | - | 1 | - | 1 | - |
| Firmicutes | *Aerococcus sanguinicola* | - | 1 | - | 1 | - | 1 | - |
| Firmicutes | *Aerococcus urinaeequi* | 2 | 2 | - | 4 | 1 | 2 | 1 |
| Firmicutes | *Aerococcus urinaehominis* | - | 3 | - | 3 | - | - | 3 |
| Firmicutes | *Aerococcus viridans* | 3 | 5 | 2 | 6 | 5 | 1 | 2 |
| Firmicutes | *Alkalihalobacillus clausii* | 5 | - | - | 5 | 5 | - | - |
| Actinobacteria | *Arcanobacterium wilhelmae* | - | 1 | 1 | - | - | 1 | - |
| Actinobacteria | *Arthrobacter gandavensis* | 1 | - | - | 1 | 1 | - | - |
| Actinobacteria | *Arthrobacter koreensis* | 1 | - | - | 1 | 1 | - | - |
| Proteobacteria | *Atlantibacter hermannii* | - | 1 | - | 1 | - | 1 | - |
| Firmicutes | *Bacillus aerius* | 3 | - | 2 | 1 | 2 | 1 | - |
| Firmicutes | *Bacillus atrophaeus* | 1 | - | - | 1 | - | 1 | - |
| Firmicutes | *Bacillus australimaris* | 1 | - | - | 1 | 1 | - | - |
| Firmicutes | *Bacillus cereus* | 1 | - | 1 | - | 1 | - | - |
| Firmicutes | *Bacillus gobiensis* | 1 | - | - | 1 | - | 1 | - |
| Firmicutes | *Bacillus haynesii* | 12 | - | 2 | 10 | 5 | 7 | - |
| Firmicutes | *Bacillus licheniformis* | 4 | 7 | 1 | 10 | 8 | 2 | 1 |
| Firmicutes | *Bacillus nakamurai* | 1 | - | - | 1 | - | 1 | - |
| Firmicutes | *Bacillus pacificus* | - | 2 | - | 2 | - | - | 2 |
| Firmicutes | *Bacillus paralicheniformis* | 30 | 20 | 9 | 41 | 19 | 20 | 11 |
| Firmicutes | *Bacillus paramobilis* | 1 | - | 1 | - | - | 1 | - |
| Firmicutes | *Bacillus piscis* | 4 | - | - | 4 | 1 | 3 | - |
| Firmicutes | *Bacillus proteolyticus* | - | 2 | - | 2 | - | 1 | 1 |
| Firmicutes | *Bacillus pumilus* | 35 | - | 15 | 20 | 27 | 8 | - |
| Firmicutes | *Bacillus sonorensis* | 2 | - | 1 | 1 | 1 | 1 | - |
| Firmicutes | *Bacillus stratosphericus* | 1 | - | - | 1 | - | 1 | - |
| Firmicutes | *Bacillus tropicus* | - | 1 | - | 1 | - | - | 1 |
| Firmicutes | *Bacillus wiedmannii* | 1 | - | 1 | - | 1 | - | - |
| Firmicutes | *Bacillus xiamenensis* | 1 | - | - | 1 | - | 1 | - |
| Firmicutes | *Bacillus zhangshouensis* | 1 | - | 1 | - | - | 1 | - |
| Firmicutes | *Brevibacillus borstelensis* | 1 | - | - | 1 | 1 | - | - |
| Firmicutes | *Caldibacillus hisashii* | - | 1 | 1 | - | - | 1 | - |
| Proteobacteria | *Campylobacter sputorum* | - | 1 | - | 1 | - | - | 1 |
| Firmicutes | *Clostridium saccharoperbutylacetonicum* | - | 1 | - | 1 | - | - | 1 |
| Actinobacteria | *Corynebacterium confusum* | 1 | - | - | 1 | 1 | - | - |
| Actinobacteria | *Corynebacterium lactis* | 3 | - | - | 3 | 3 | - | - |
| Actinobacteria | *Corynebacterium renale* | 11 | 16 | 9 | 18 | 15 | 1 | 11 |
| Actinobacteria | *Corynebacterium vitaeruminis* | 1 | - | - | 1 | 1 | - | - |
| Actinobacteria | *Corynebacterium xerosis* | 2 | - | - | 2 | 2 | - | - |

**Supplemental Table 1 Cont’d.** A list of all bacterial isolates recovered, their growth media and oxygen condition, and their anatomic origin.
| Phylum | Bacteria Name | Number of Isolates |  | Isolate Origin |  | Media |  |  |
| --- | --- | --- | --- | --- | --- | --- | --- | --- |
|  |  | Aerobic | Anaerobic | Uterus | Vagina | CB/Blood | MRS | WC |
| Deferribacteres | *Denitrovibrio acetiphilus* | - | 1 | - | 1 | - | - | 1 |
| Firmicutes | *Enterococcus aquimarinus* | - | 1 | 1 | - | 1 | - | - |
| Firmicutes | *Enterococcus casseliflavus* | - | 1 | - | 1 | - | - | 1 |
| Firmicutes | *Enterococcus gallinarum* | - | 3 | - | 3 | 1 | 2 | - |
| Firmicutes | *Enterococcus hirae* | - | 2 | 1 | 1 | - | 2 | - |
| Firmicutes | *Erysipelothrix inopinata* | - | 1 | 1 | - | 1 | - | - |
| Proteobacteria | *Escherichia coli* | - | 1 | - | 1 | - | 1 | - |
| Proteobacteria | *Escherichia fergusonii* | - | 11 | 4 | 7 | 1 | 6 | 4 |
| Proteobacteria | *Escherichia fergusonii* ATCC 35469 | 25 | 1 | 9 | 17 | 16 | 10 | - |
| Proteobacteria | *Escherichia marmotae* | 9 | 1 | 1 | 9 | 6 | 4 | - |
| Firmicutes | *Facklamia hominis* | 4 | 8 | 6 | 6 | 8 | 3 | 1 |
| Proteobacteria | *Gallibacterium salpingitidis* | 2 | 2 | 4 | - | 2 | 1 | 1 |
| Firmicutes | *Helcococcus ovis* | 1 | - | - | 1 | 1 | - | - |
| Firmicutes | *Heyndrickxia oleronia* | 5 | - | - | 5 | 1 | 4 | - |
| Proteobacteria | *Histophilus somni* | 2 | - | 1 | 1 | 2 | - | - |
| Firmicutes | *Hutsoniella sourekii* | 2 | 4 | - | 6 | 3 | - | 3 |
| Proteobacteria | *Hydrogenophaga electricum* | 1 | - | - | 1 | 1 | - | - |
| Proteobacteria | *Kosakonia oryziphila* | 1 | - | - | 1 | - | 1 | - |
| Actinobacteria | *Kribbella monticola* | - | 1 | 1 | - | 1 | - | - |
| Firmicutes | *Kurthia gibsonii* | - | 1 | - | 1 | - | - | 1 |
| Firmicutes | *Lacrimispora algidixylanolytica* | - | 1 | - | 1 | - | 1 | - |
| Firmicutes | *Lactacaseibacillus pantheris* | 1 | - | 1 | - | 1 | - | - |
| Firmicutes | *Lactiplantibacillus plantarum* | - | 1 | - | 1 | - | 1 | - |
| Firmicutes | *Lactobacillus amylovorus* | - | 2 | - | 2 | 1 | 1 | - |
| Proteobacteria | *Lelliottia amnigena* | - | 1 | - | 1 | - | - | 1 |
| Firmicutes | *Limosilactobacillus mucosae* | - | 5 | 4 | 1 | 1 | 4 | - |
| Firmicutes | *Lysinibacillus macroides* | 2 | - | - | 2 | 1 | 1 | - |
| Firmicutes | *Lysinibacillus pakistanensis* | 1 | - | - | 1 | - | 1 | - |
| Firmicutes | *Macrococcus canis* | 4 | - | 1 | 3 | 4 | - | - |
| Proteobacteria | *Mannheimia granulomatis* | - | 1 | - | 1 | - | - | 1 |
| Proteobacteria | *Mannheimia ruminalis* | 2 | - | - | 2 | 2 | - | - |
| Proteobacteria | *Mannheimia varigena* | 1 | 3 | 1 | 3 | 1 | - | 3 |
| Proteobacteria | *Moraxella bovis* | 1 | - | - | 1 | 1 | - | - |
| Proteobacteria | *Moraxella bovoculi* | 2 | - | 2 | - | 2 | - | - |
| Proteobacteria | *Moraxella catarrhalis* | 1 | - | - | 1 | 1 | - | - |
| Proteobacteria | *Moraxella osloensis* | 3 | - | 2 | 1 | 3 | - | - |
| Proteobacteria | *Neisseria weaveri* | - | 1 | 1 | - | - | - | 1 |
| Firmicutes | *Neobacillus cucumis* | - | 1 | - | 1 | 1 | - | - |
| Firmicutes | *Niallia circulans* | 3 | 1 | 1 | 3 | 3 | 1 | - |
| Firmicutes | *Paenibacillus borealis* | - | 1 | - | 1 | - | - | 1 |
| Firmicutes | *Paenibacillus campinasensis* | 1 | - | - | 1 | 1 | - | - |
| Firmicutes | *Paenibacillus daejeonensis* | 1 | - | 1 | - | 1 | - | - |
| Firmicutes | *Paenibacillus lautus* | 1 | - | - | 1 | 1 | - | - |

**Supplemental Table 1 Cont'd.** A list of all bacterial isolates recovered, their growth media and oxygen condition, and their anatomic origin.
| Phylum | Bacteria Name | Number of Isolates |  | Isolate Origin |  | Media |  |  |
| --- | --- | --- | --- | --- | --- | --- | --- | --- |
|  |  | Aerobic | Anaerobic | Uterus | Vagina | CB/Blood | MRS | WC |
| Firmicutes | *Paenibacillus macerans* | - | 1 | - | 1 | - | - | 1 |
| Firmicutes | *Paenibacillus mobilis* | - | 1 | - | 1 | - | - | 1 |
| Firmicutes | *Paenibacillus naphthalenovorans* | 5 | - | - | 5 | 5 | - | - |
| Firmicutes | *Paenibacillus polysaccharolyticus* | - | 1 | - | 1 | 1 | - | - |
| Firmicutes | *Paenibacillus silvae* | 1 | - | - | 1 | 1 | - | - |
| Proteobacteria | *Pelistega europaea* | 1 | - | 1 | - | 1 | - | - |
| Firmicutes | *Peptoniphilus indolicus* | - | 7 | - | 7 | 4 | 1 | 2 |
| Firmicutes | *Peribacillus psychrosaccharolyticus* | - | 1 | - | 1 | - | - | 1 |
| Firmicutes | *Periweissella fabalis* | - | 1 | - | 1 | - | - | 1 |
| Firmicutes | *Priestia aryabhatai* | 1 | - | 1 | - | - | 1 | - |
| Firmicutes | *Priestia megaterium* | 2 | - | 1 | 1 | 1 | 1 | - |
| Proteobacteria | *Proteus mirabilis* | 1 | 9 | - | 10 | 6 | 1 | 3 |
| Proteobacteria | *Pseudescerichia vulneris* | 1 | - | - | 1 | - | 1 | - |
| Proteobacteria | *Pseudomonas flexibilis* | 2 | - | 2 | - | 2 | - | - |
| Firmicutes | *Robertmurraya siralis* | 1 | - | - | 1 | 1 | - | - |
| Firmicutes | *Rossellomorea marisflavi* | 1 | - | - | 1 | - | 1 | - |
| Actinobacteria | *Rothia nasimurium* | 1 | - | - | 1 | 1 | - | - |
| Proteobacteria | *Shigella boydii* | 1 | - | - | 1 | 1 | - | - |
| Proteobacteria | *Shigella flexneri* | 1 | 1 | 1 | 1 | 1 | 1 | = |
| Proteobacteria | *Shigella sonnei* | - | 1 | 1 | - | - | - | 1 |
| Firmicutes | *Solibacillus isronensis* | 1 | - | 1 | - | 1 | - | - |
| Firmicutes | *Staphylococcus aureus* | - | 1 | - | 1 | - | 1 | - |
| Firmicutes | *Staphylococcus capitis* | 2 | - | 2 | - | 2 | - | - |
| Firmicutes | *Staphylococcus chromogenes* | 6 | 11 | 2 | 15 | 9 | 1 | 7 |
| Firmicutes | *Staphylococcus epidermidis* | 1 | 1 | - | 2 | 2 | - | - |
| Firmicutes | *Staphylococcus hominis* | - | 1 | - | 1 | 1 | - | - |
| Firmicutes | *Staphylococcus hyicus* | 6 | - | 1 | 5 | 5 | 1 | - |
| Firmicutes | *Staphylococcus pasteurii* | - | 1 | - | 1 | - | 1 | - |
| Firmicutes | *Staphylococcus roterodami* | 1 | - | - | 1 | 1 | - | - |
| Firmicutes | *Streptococcus anginosus* | - | 1 | - | 1 | - | - | 1 |
| Firmicutes | *Streptococcus equinus* | 2 | 38 | 9 | 31 | 13 | 16 | 11 |
| Firmicutes | *Streptococcus equinus* ATCC 9812 | 8 | - | 7 | 1 | 2 | 6 | - |
| Firmicutes | *Streptococcus lutetiensis* | 4 | 5 | 2 | 7 | 3 | 5 | 1 |
| Firmicutes | *Streptococcus pluranimalium* | 82 | 151 | 88 | 145 | 117 | 83 | 33 |
| Firmicutes | *Streptococcus ruminantium* | 1 | - | - | 1 | 1 | - | - |
| Firmicutes | *Streptococcus suis* | 1 | 3 | 1 | 3 | 1 | - | 3 |
| Actinobacteria | *Streptomyces viridodiiastaticus* | 1 | - | - | 1 | 1 | - | - |
| Proteobacteria | *Thioalkalivibrio jannaschii* | - | 1 | - | 1 | - | - | 1 |
| Actinobacteria | *Trueperella abortus suis* | 12 | - | 1 | 11 | 12 | - | - |
| Firmicutes | *Weizmannia coagulans* | - | 2 | - | 2 | - | 2 | - |
| Total number of isolates |  | 363 | 370 | 216 | 517 | 377 | 226 | 130 |

**Supplementary Table 2.**
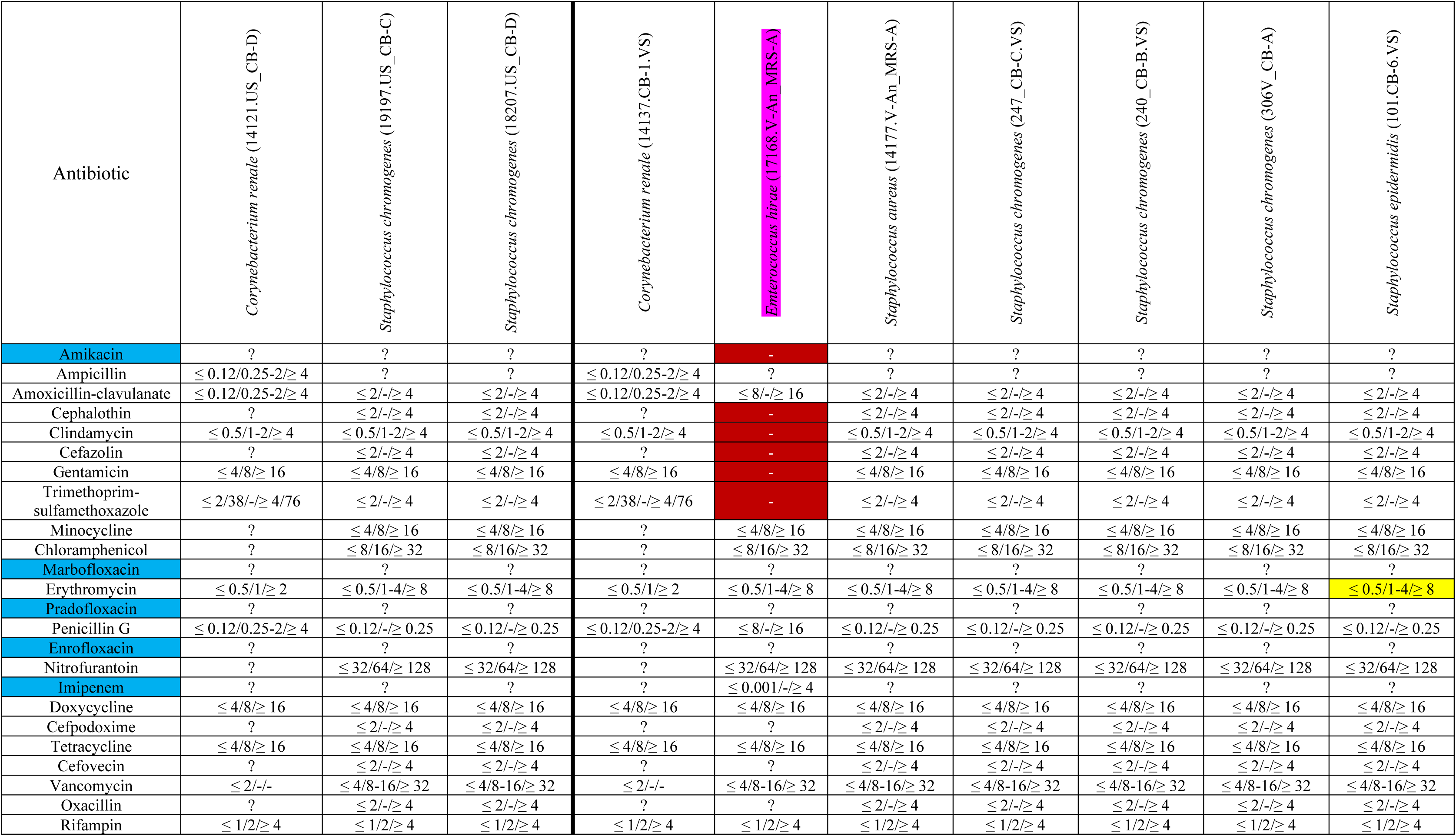
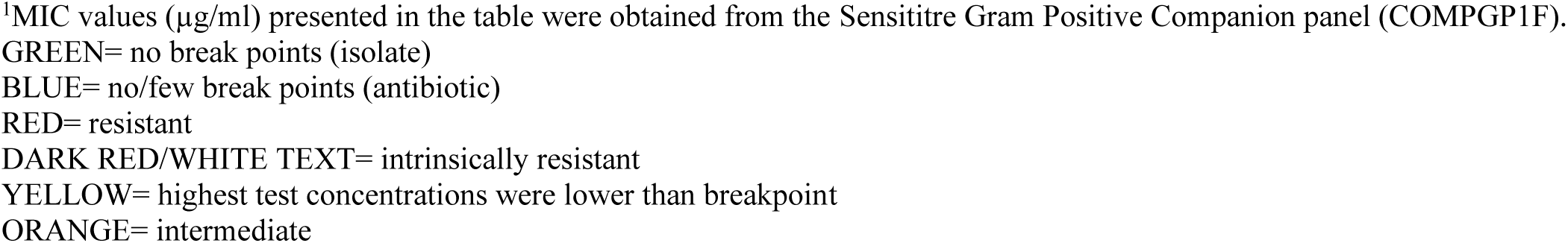
Antimicrobial resistance breakpoints that have been reported previously (Gram-positive) (CLSI, 2016, 2021, 2022).^1^.

| Antibiotic | *Corynebacterium renale* (14121.US_CB-D) | *Staphylococcus chromogenes* (19197.US_CB-C) | *Staphylococcus chromogenes* (18207.US_CB-D) | *Corynebacterium renale* (14137.CB-1.VS) | *Enterococcus hirae* (17168.V-An_MRS-A) | *Staphylococcus aureus* (14177.V-An_MRS-A) | *Staphylococcus chromogenes* (247_CB-C.VS) | *Staphylococcus chromogenes* (240_CB-B.VS) | *Staphylococcus chromogenes* (306V_CB-A) | *Staphylococcus epidermidis* (101.CB-6.VS) |
| --- | --- | --- | --- | --- | --- | --- | --- | --- | --- | --- |
| Amikacin | ? | ? | ? | ? | - | ? | ? | ? | ? | ? |
| Ampicillin | $\leq 0.12/0.25-2/\geq 4$ | ? | ? | $\leq 0.12/0.25-2/\geq 4$ | ? | ? | ? | ? | ? | ? |
| Amoxicillin-clavulanate | $\leq 0.12/0.25-2/\geq 4$ | $\leq 2/-/\geq 4$ | $\leq 2/-/\geq 4$ | $\leq 0.12/0.25-2/\geq 4$ | $\leq 8/-/\geq 16$ | $\leq 2/-/\geq 4$ | $\leq 2/-/\geq 4$ | $\leq 2/-/\geq 4$ | $\leq 2/-/\geq 4$ | $\leq 2/-/\geq 4$ |
| Cephalothin | ? | $\leq 2/-/\geq 4$ | $\leq 2/-/\geq 4$ | ? | - | $\leq 2/-/\geq 4$ | $\leq 2/-/\geq 4$ | $\leq 2/-/\geq 4$ | $\leq 2/-/\geq 4$ | $\leq 2/-/\geq 4$ |
| Clindamycin | $\leq 0.5/1-2/\geq 4$ | $\leq 0.5/1-2/\geq 4$ | $\leq 0.5/1-2/\geq 4$ | $\leq 0.5/1-2/\geq 4$ | - | $\leq 0.5/1-2/\geq 4$ | $\leq 0.5/1-2/\geq 4$ | $\leq 0.5/1-2/\geq 4$ | $\leq 0.5/1-2/\geq 4$ | $\leq 0.5/1-2/\geq 4$ |
| Cefazolin | ? | $\leq 2/-/\geq 4$ | $\leq 2/-/\geq 4$ | ? | - | $\leq 2/-/\geq 4$ | $\leq 2/-/\geq 4$ | $\leq 2/-/\geq 4$ | $\leq 2/-/\geq 4$ | $\leq 2/-/\geq 4$ |
| Gentamicin | $\leq 4/8/\geq 16$ | $\leq 4/8/\geq 16$ | $\leq 4/8/\geq 16$ | $\leq 4/8/\geq 16$ | - | $\leq 4/8/\geq 16$ | $\leq 4/8/\geq 16$ | $\leq 4/8/\geq 16$ | $\leq 4/8/\geq 16$ | $\leq 4/8/\geq 16$ |
| Trimethoprim-sulfamethoxazole | $\leq 2/38/-/\geq 4/76$ | $\leq 2/-/\geq 4$ | $\leq 2/-/\geq 4$ | $\leq 2/38/-/\geq 4/76$ | - | $\leq 2/-/\geq 4$ | $\leq 2/-/\geq 4$ | $\leq 2/-/\geq 4$ | $\leq 2/-/\geq 4$ | $\leq 2/-/\geq 4$ |
| Minocycline | ? | $\leq 4/8/\geq 16$ | $\leq 4/8/\geq 16$ | ? | $\leq 4/8/\geq 16$ | $\leq 4/8/\geq 16$ | $\leq 4/8/\geq 16$ | $\leq 4/8/\geq 16$ | $\leq 4/8/\geq 16$ | $\leq 4/8/\geq 16$ |
| Chloramphenicol | ? | $\leq 8/16/\geq 32$ | $\leq 8/16/\geq 32$ | ? | $\leq 8/16/\geq 32$ | $\leq 8/16/\geq 32$ | $\leq 8/16/\geq 32$ | $\leq 8/16/\geq 32$ | $\leq 8/16/\geq 32$ | $\leq 8/16/\geq 32$ |
| Marbofloxacin | ? | ? | ? | ? | ? | ? | ? | ? | ? | ? |
| Erythromycin | $\leq 0.5/1/\geq 2$ | $\leq 0.5/1-4/\geq 8$ | $\leq 0.5/1-4/\geq 8$ | $\leq 0.5/1/\geq 2$ | $\leq 0.5/1-4/\geq 8$ | $\leq 0.5/1-4/\geq 8$ | $\leq 0.5/1-4/\geq 8$ | $\leq 0.5/1-4/\geq 8$ | $\leq 0.5/1-4/\geq 8$ | $\leq 0.5/1-4/\geq 8$ |
| Pradofloxacin | ? | ? | ? | ? | ? | ? | ? | ? | ? | ? |
| Penicillin G | $\leq 0.12/0.25-2/\geq 4$ | $\leq 0.12/-/\geq 0.25$ | $\leq 0.12/-/\geq 0.25$ | $\leq 0.12/0.25-2/\geq 4$ | $\leq 8/-/\geq 16$ | $\leq 0.12/-/\geq 0.25$ | $\leq 0.12/-/\geq 0.25$ | $\leq 0.12/-/\geq 0.25$ | $\leq 0.12/-/\geq 0.25$ | $\leq 0.12/-/\geq 0.25$ |
| Enrofloxacin | ? | ? | ? | ? | ? | ? | ? | ? | ? | ? |
| Nitrofurantoin | ? | $\leq 32/64/\geq 128$ | $\leq 32/64/\geq 128$ | ? | $\leq 32/64/\geq 128$ | $\leq 32/64/\geq 128$ | $\leq 32/64/\geq 128$ | $\leq 32/64/\geq 128$ | $\leq 32/64/\geq 128$ | $\leq 32/64/\geq 128$ |
| Imipenem | ? | ? | ? | ? | $\leq 0.001/-/\geq 4$ | ? | ? | ? | ? | ? |
| Doxycycline | $\leq 4/8/\geq 16$ | $\leq 4/8/\geq 16$ | $\leq 4/8/\geq 16$ | $\leq 4/8/\geq 16$ | $\leq 4/8/\geq 16$ | $\leq 4/8/\geq 16$ | $\leq 4/8/\geq 16$ | $\leq 4/8/\geq 16$ | $\leq 4/8/\geq 16$ | $\leq 4/8/\geq 16$ |
| Cefpodoxime | ? | $\leq 2/-/\geq 4$ | $\leq 2/-/\geq 4$ | ? | ? | $\leq 2/-/\geq 4$ | $\leq 2/-/\geq 4$ | $\leq 2/-/\geq 4$ | $\leq 2/-/\geq 4$ | $\leq 2/-/\geq 4$ |
| Tetracycline | $\leq 4/8/\geq 16$ | $\leq 4/8/\geq 16$ | $\leq 4/8/\geq 16$ | $\leq 4/8/\geq 16$ | $\leq 4/8/\geq 16$ | $\leq 4/8/\geq 16$ | $\leq 4/8/\geq 16$ | $\leq 4/8/\geq 16$ | $\leq 4/8/\geq 16$ | $\leq 4/8/\geq 16$ |
| Cefovecin | ? | $\leq 2/-/\geq 4$ | $\leq 2/-/\geq 4$ | ? | ? | $\leq 2/-/\geq 4$ | $\leq 2/-/\geq 4$ | $\leq 2/-/\geq 4$ | $\leq 2/-/\geq 4$ | $\leq 2/-/\geq 4$ |
| Vancomycin | $\leq 2/-/-$ | $\leq 4/8-16/\geq 32$ | $\leq 4/8-16/\geq 32$ | $\leq 2/-/-$ | $\leq 4/8-16/\geq 32$ | $\leq 4/8-16/\geq 32$ | $\leq 4/8-16/\geq 32$ | $\leq 4/8-16/\geq 32$ | $\leq 4/8-16/\geq 32$ | $\leq 4/8-16/\geq 32$ |
| Oxacillin | ? | $\leq 2/-/\geq 4$ | $\leq 2/-/\geq 4$ | ? | ? | $\leq 2/-/\geq 4$ | $\leq 2/-/\geq 4$ | $\leq 2/-/\geq 4$ | $\leq 2/-/\geq 4$ | $\leq 2/-/\geq 4$ |
| Rifampin | $\leq 1/2/\geq 4$ | $\leq 1/2/\geq 4$ | $\leq 1/2/\geq 4$ | $\leq 1/2/\geq 4$ | $\leq 1/2/\geq 4$ | $\leq 1/2/\geq 4$ | $\leq 1/2/\geq 4$ | $\leq 1/2/\geq 4$ | $\leq 1/2/\geq 4$ | $\leq 1/2/\geq 4$ |

<sup>1</sup>MIC values (µg/ml) presented in the table were obtained from the Sensititre Gram Positive Companion panel (COMPGP1F). GREEN= no break points (isolate) BLUE= no/few break points (antibiotic) RED= resistant DARK RED/WHITE TEXT= intrinsically resistant YELLOW= highest test concentrations were lower than breakpoint ORANGE= intermediate

**Supplementary Table 3.** Antimicrobial resistance breakpoints that have been reported previously (Gram-negative) (Goldspink et al., 2015; CLSI, 2016; Gomes et al., 2019; CLSI, 2021, 2022)^1^.

| Antibiotic | *Escherichia coli* (18207.US_CB-C) | *Escherichia coli* (14121.US_MRS-B) | *Escherichia coli* (18333.US_CB-C) | *Histophilus somni* (20116.US_CB-A) | *Actinobacillus seminis* (306V_CB-B) | *Escherichia coli* (169_CB-A.VS) | *Escherichia coli* (7091.V-An_Bld-A) | *Escherichia coli* (18081.CB-1.VS) | *Escherichia coli* (247_CB-D.VS) | *Escherichia coli* (247_MRS-A.VS) |
| --- | --- | --- | --- | --- | --- | --- | --- | --- | --- | --- |
| Amikacin | ≤ 16/32/≥ 64 | ≤ 16/32/≥ 64 | ≤ 16/32/≥ 64 | - | - | ≤ 16/32/≥ 64 | ≤ 16/32/≥ 64 | ≤ 16/32/≥ 64 | ≤ 16/32/≥ 64 | ≤ 16/32/≥ 64 |
| Ampicillin | ≤ 8/16/≥ 32 | ≤ 8/16/≥ 32 | ≤ 8/16/≥ 32 | ? | ? | ≤ 8/16/≥ 32 | ≤ 8/16/≥ 32 | ≤ 8/16/≥ 32 | ≤ 8/16/≥ 32 | ≤ 8/16/≥ 32 |
| Azithromycin | ≤ 16/-/≥ 32 | ≤ 16/-/≥ 32 | ≤ 16/-/≥ 32 | - | - | ≤ 16/-/≥ 32 | ≤ 16/-/≥ 32 | ≤ 16/-/≥ 32 | ≤ 16/-/≥ 32 | ≤ 16/-/≥ 32 |
| Cefitofur | - | - | - | ≤ 2/-/≥ 8 | ? | - | - | - | - | - |
| Clindamycin | - | - | - | ? | ? | - | - | - | - | - |
| Danofloxacin | - | - | - | ? | ? | - | - | - | - | - |
| Cefazolin | ≤ 2/4/≥ 8 | ≤ 2/4/≥ 8 | ≤ 2/4/≥ 8 | - | - | ≤ 2/4/≥ 8 | ≤ 2/4/≥ 8 | ≤ 2/4/≥ 8 | ≤ 2/4/≥ 8 | ≤ 2/4/≥ 8 |
| Gentamicin | ≤ 4/8/≥ 16 | ≤ 4/8/≥ 16 | ≤ 4/8/≥ 16 | ? | ? | ≤ 4/8/≥ 16 | ≤ 4/8/≥ 16 | ≤ 4/8/≥ 16 | ≤ 4/8/≥ 16 | ≤ 4/8/≥ 16 |
| Gamithromycin | - | - | - | ? | ? | - | - | - | - | - |
| Trimethoprim-sulfamethoxazole | ≤ 2/-/≥ 4 | ≤ 2/-/≥ 4 | ≤ 2/-/≥ 4 | ? | ? | ≤ 2/-/≥ 4 | ≤ 2/-/≥ 4 | ≤ 2/-/≥ 4 | ≤ 2/-/≥ 4 | ≤ 2/-/≥ 4 |
| Ceftiofur | ≤ 2/4/≥ 8 | ≤ 2/4/≥ 8 | ≤ 2/4/≥ 8 | ? | - | ≤ 2/4/≥ 8 | ≤ 2/4/≥ 8 | ≤ 2/4/≥ 8 | ≤ 2/4/≥ 8 | ≤ 2/4/≥ 8 |
| Chloramphenicol | ≤ 8/16/≥ 32 | ≤ 8/16/≥ 32 | ≤ 8/16/≥ 32 | - | - | ≤ 8/16/≥ 32 | ≤ 8/16/≥ 32 | ≤ 8/16/≥ 32 | ≤ 8/16/≥ 32 | ≤ 8/16/≥ 32 |
| Ceftazidime | ≤ 4/8/≥ 16 | ≤ 4/8/≥ 16 | ≤ 4/8/≥ 16 | - | - | ≤ 4/8/≥ 16 | ≤ 4/8/≥ 16 | ≤ 4/8/≥ 16 | ≤ 4/8/≥ 16 | ≤ 4/8/≥ 16 |
| Erythromycin | ? | ? | ? | - | - | ? | ? | ? | ? | ? |
| Clarithromycin | ? | ? | ? | - | - | ? | ? | ? | ? | ? |
| Penicillin G | ? | ? | ? | ? | ? | ? | ? | ? | ? | ? |
| Enrofloxacin | ? | ? | ? | ≤ 0.25/-/≥ 2 | ? | ? | ? | ? | ? | ? |
| Imipenem | ≤ 1/2/≥ 4 | ≤ 1/2/≥ 4 | ≤ 1/2/≥ 4 | - | - | ≤ 1/2/≥ 4 | ≤ 1/2/≥ 4 | ≤ 1/2/≥ 4 | ≤ 1/2/≥ 4 | ≤ 1/2/≥ 4 |
| Doxycycline | ≤ 4/8/≥ 16 | ≤ 4/8/≥ 16 | ≤ 4/8/≥ 16 | - | - | ≤ 4/8/≥ 16 | ≤ 4/8/≥ 16 | ≤ 4/8/≥ 16 | ≤ 4/8/≥ 16 | ≤ 4/8/≥ 16 |
| Tetracycline | ≤ 4/8/≥ 16 | ≤ 4/8/≥ 16 | ≤ 4/8/≥ 16 | ≤ 2/-/≥ 8 | ? | ≤ 4/8/≥ 16 | ≤ 4/8/≥ 16 | ≤ 4/8/≥ 16 | ≤ 4/8/≥ 16 | ≤ 4/8/≥ 16 |
| Ticarcillin | ≤ 16/32-64/≥ 128 | ≤ 16/32-64/≥ 128 | ≤ 16/32-64/≥ 128 | - | - | ≤ 16/32-64/≥ 128 | ≤ 16/32-64/≥ 128 | ≤ 16/32-64/≥ 128 | ≤ 16/32-64/≥ 128 | ≤ 16/32-64/≥ 128 |
| Ticarcillin-clavulanate | ? | ? | ? | - | - | ? | ? | ? | ? | ? |
| Oxacillin | ? | ? | ? | - | - | ? | ? | ? | ? | ? |
| Rifampin | ? | ? | ? | - | - | ? | ? | ? | ? | ? |
| Sulfadimethoxine | - | - | - | ? | ? | - | - | - | - | - |
| Tiamulin | - | - | - | ? | ? | - | - | - | - | - |
| Tildipirosin | - | - | - | ? | ? | - | - | - | - | - |
| Tilmicosin | - | - | - | $\leq 8/-/\geq 32$ | ? | - | - | - | - | - |
| Florfenicol | - | - | - | $\leq 16/-/\geq 64$ | ? | - | - | - | - | - |
| Tulathromycin | - | - | - | ? | ? | - | - | - | - | - |
| Tylosin tartrate | - | - | - | ? | ? | - | - | - | - | - |
| Neomycin | - | - | - | ? | ? | - | - | - | - | - |
| Spectinomycin | - | - | - | ? | - | - | - | - | - | - |

**Supplementary Table 3 Cont’d.** Antimicrobial resistance breakpoints that have been reported previously (Gram-negative) (CLSI, 2016; Gomes et al., 2019; CLSI, 2021, 2022)<sup>1</sup>.
| Antibiotic | *Escherichia coli* (96_MRS-B.VS) | *Escherichia coli* (240_CB-A.VS) | *Escherichia coli* ( 46_CB-A.VS) | *Escherichia coli* (240_MRS-A.VS) | *Escherichia coli* (247_CB-B.VS) | *Escherichia coli* (267_CB-A.VS) | *Escherichia coli* (19044.V-An_MRS-A) | *Escherichia coli* (247_MRS-C.VS) | *Histophilus somni* (029V_CB-B) |
| --- | --- | --- | --- | --- | --- | --- | --- | --- | --- |
| Amikacin | ≤ 16/32/≥ 64 | ≤ 16/32/≥ 64 | ≤ 16/32/≥ 64 | ≤ 16/32/≥ 64 | ≤ 16/32/≥ 64 | ≤ 16/32/≥ 64 | ≤ 16/32/≥ 64 | ≤ 16/32/≥ 64 | - |
| Ampicillin | ≤ 8/16/≥ 32 | ≤ 8/16/≥ 32 | ≤ 8/16/≥ 32 | ≤ 8/16/≥ 32 | ≤ 8/16/≥ 32 | ≤ 8/16/≥ 32 | ≤ 8/16/≥ 32 | ≤ 8/16/≥ 32 | ? |
| Azithromycin | ≤ 16/-/≥ 32 | ≤ 16/-/≥ 32 | ≤ 16/-/≥ 32 | ≤ 16/-/≥ 32 | ≤ 16/-/≥ 32 | ≤ 16/-/≥ 32 | ≤ 16/-/≥ 32 | - | - |
| Cefitofur | - | - | - | - | - | - | - | - | ≤ 2/-/≥ 8 |
| Clindamycin | - | - | - | - | - | - | - | ? | ? |
| Danofloxacin | - | - | - | - | - | - | - | - | ? |
| Cefazolin | ≤ 2/4/≥ 8 | ≤ 2/4/≥ 8 | ≤ 2/4/≥ 8 | ≤ 2/4/≥ 8 | ≤ 2/4/≥ 8 | ≤ 2/4/≥ 8 | ≤ 2/4/≥ 8 | ≤ 2/4/≥ 8 | - |
| Gentamicin | ≤ 4/8/≥ 16 | ≤ 4/8/≥ 16 | ≤ 4/8/≥ 16 | ≤ 4/8/≥ 16 | ≤ 4/8/≥ 16 | ≤ 4/8/≥ 16 | ≤ 4/8/≥ 16 | ≤ 4/8/≥ 16 | ? |
| Gamithromycin | - | - | - | - | - | - | - | - | ? |
| Trimethoprim-sulfamethoxazole | ≤ 2/-/≥ 4 | ≤ 2/-/≥ 4 | ≤ 2/-/≥ 4 | ≤ 2/-/≥ 4 | ≤ 2/-/≥ 4 | ≤ 2/-/≥ 4 | ≤ 2/-/≥ 4 | ≤ 2/-/≥ 4 | ? |
| Ceftiofur | ≤ 2/4/≥ 8 | ≤ 2/4/≥ 8 | ≤ 2/4/≥ 8 | ≤ 2/4/≥ 8 | ≤ 2/4/≥ 8 | ≤ 2/4/≥ 8 | ≤ 2/4/≥ 8 | - | ? |
| Chloramphenicol | ≤ 8/16/≥ 32 | ≤ 8/16/≥ 32 | ≤ 8/16/≥ 32 | ≤ 8/16/≥ 32 | ≤ 8/16/≥ 32 | ≤ 8/16/≥ 32 | ≤ 8/16/≥ 32 | ≤ 8/16/≥ 32 | - |
| Ceftazidime | ≤ 4/8/≥ 16 | ≤ 4/8/≥ 16 | ≤ 4/8/≥ 16 | ≤ 4/8/≥ 16 | ≤ 4/8/≥ 16 | ≤ 4/8/≥ 16 | ≤ 4/8/≥ 16 | - | - |
| Erythromycin | ? | ? | ? | ? | ? | ? | ? | ? | - |
| Clarithromycin | ? | ? | ? | ? | ? | ? | ? | - | - |
| Penicillin G | ? | ? | ? | ? | ? | ? | ? | ? | ? |
| Enrofloxacin | ? | ? | ? | ? | ? | ? | ? | ? | ≤ 0.25/-/≥ 2 |
| Imipenem | ≤ 1/2/≥ 4 | ≤ 1/2/≥ 4 | ≤ 1/2/≥ 4 | ≤ 1/2/≥ 4 | ≤ 1/2/≥ 4 | ≤ 1/2/≥ 4 | ≤ 1/2/≥ 4 | ≤ 1/2/≥ 4 | - |
| Doxycycline | ≤ 4/8/≥ 16 | ≤ 4/8/≥ 16 | ≤ 4/8/≥ 16 | ≤ 4/8/≥ 16 | ≤ 4/8/≥ 16 | ≤ 4/8/≥ 16 | ≤ 4/8/≥ 16 | ≤ 4/8/≥ 16 | - |
| Tetracycline | ≤ 4/8/≥ 16 | ≤ 4/8/≥ 16 | ≤ 4/8/≥ 16 | ≤ 4/8/≥ 16 | ≤ 4/8/≥ 16 | ≤ 4/8/≥ 16 | ≤ 4/8/≥ 16 | ≤ 4/8/≥ 16 | ≤ 2/-/≥ 8 |
| Ticarcillin | ≤ 16/32-64/≥ 128 | ≤ 16/32-64/≥ 128 | ≤ 16/32-64/≥ 128 | ≤ 16/32-64/≥ 128 | ≤ 16/32-64/≥ 128 | ≤ 16/32-64/≥ 128 | ≤ 16/32-64/≥ 128 | - | - |
| Ticarcillin-clavulanate | ? | ? | ? | ? | ? | ? | ? | - | - |
| Oxacillin | ? | ? | ? | ? | ? | ? | ? | ? | - |
| Rifampin | ? | ? | ? | ? | ? | ? | ? | ? | - |
| Sulfadimethoxine | - | - | - | - | - | - | - | - | ? |
| Tiamulin | - | - | - | - | - | - | - | - | ? |
| Tildipirosin | - | - | - | - | - | - | - | - | ? |
| Tilmicosin | - | - | - | - | - | - | - | - | $\leq 8/-\geq 32$ |
| Florfenicol | - | - | - | - | - | - | - | - | $\leq 16/-\geq 64$ |
| Tulathromycin | - | - | - | - | - | - | - | - | ? |
| Tylosin tartrate | - | - | - | - | - | - | - | - | ? |
| Neomycin | - | - | - | - | - | - | - | - | ? |
| Spectinomycin | - | - | - | - | - | - | - | - | ? |

<sup>1</sup>MIC values (µg/ml) presented in the table were obtained from the Sensititre Gram Negative Companion panel (COMPGN1F).
GREEN= no/few break points (isolate)
BLUE= no/few break points (antibiotic)
RED= resistant
DARK RED/WHITE TEXT= intrinsically resistant
YELLOW= highest test concentrations were lower than breakpoints
ORANGE= intermediate

## Notes

### Competing Interest Statement

The authors have declared no competing interest.

